# Diagnosing and predicting mixed culture fermentations with unicellular and guild-based metabolic models

**DOI:** 10.1101/759548

**Authors:** Matthew J. Scarborough, Joshua J. Hamilton, Elizabeth A. Erb, Timothy J. Donohue, Daniel R. Noguera

**Affiliations:** The Great Lakes Bioenergy Research Center, UW-Madison, Madison, WI; Department of Civil and Environmental Engineering, UW-Madison, Madison, WI; Department of Civil and Environmental Engineering, University of Vermont, Burlington, VT; Department of Biochemistry, UW-Madison, Madison, WI; Department of Chemical and Biological Engineering, UW-Madison, Madison, WI; Department of Bacteriology, UW-Madison, Madison, WI

## Abstract

Multi-species microbial communities determine the fate of materials in the environment and can be harnessed to produce beneficial products from renewable resources. In a recent example, fermentations by microbial communities have produced medium-chain fatty acids (MCFAs). Tools to predict, assess, and improve the performance of these communities, however, are limited. To provide such tools, we constructed two metabolic models of MCFA-producing microbial communities based on available genomic, transcriptomic and metabolomic data. The first model is a unicellular model (iFermCell215), while the second model (iFermGuilds789) separates fermentation activities into functional guilds. Ethanol and lactate are fermentation products known to serve as substrates for MCFA production, while acetate is another common co-metabolite during MCFA production. Simulations with iFermCell215 predict that low molar ratios of acetate-to-ethanol favor MCFA production, whereas the products of lactate and acetate co-utilization are less dependent on the acetate-to-lactate ratio. In simulations of an MCFA-producing community fed a complex organic mixture derived from lignocellulose, iFermGuilds789 predicted that lactate was an extracellular co-metabolite that served as a substrate for butyrate (C4) production. Extracellular hexanoic (C6) and octanoic acids (C8) were predicted by iFermGuilds789 to be from community members that directly metabolize sugars. Modeling results provide several hypotheses that can improve our understanding of microbial roles in a MCFA-producing microbiome and inform strategies to increase MCFA production. Further, these models represent novel tools for exploring the role of mixed microbial communities in carbon recycling in the environment, as well as on beneficial reuse of organic residues.

**IMPORTANCE:** Microbiomes are vital to human health, agriculture, and protecting the environment. Predicting behavior of self-assembled or synthetic microbiomes, however, remains a challenge. In this work, we used unicellular and guild-based metabolic models to investigate production of medium-chain fatty acids by a mixed microbial community that is fed multiple organic substrates. Modeling results provided insights into metabolic pathways of three medium-chain fatty acid-producing guilds and identified potential strategies to increase production of medium-chain fatty acids. This work demonstrates the role of metabolic models in augmenting multi-omic studies to gain greater insights into microbiome behavior.

## INTRODUCTION

Mixed microbial fermentations have benefited humanity throughout history.^1^ Since the mid-18^th^ century, mixed culture fermentations have produced valuable chemicals and recovered energy from wastes.^2^ Many mixed microbial fermentations involve the synergistic activities of several functional groups to convert organic substrates into valuable products. In anaerobic digestion, fermentation products like acetic acid (C2) and H_2_ are used to support microbial production of methane, thus enabling the recovery of a large fraction of the chemical energy originally present in the complex organic substrates. While anaerobic digestion technologies are well established, other strategies to expand the range of potential products of mixed culture fermentations are only beginning to emerge. One example of such a mixed culture fermentation is the carboxylate platform, which produces medium-chain fatty acids (MCFAs) as a potentially valuable set of bio-based products.^3^

MCFAs are attractive carboxylate platform products because of their many industrial uses.^4^ Octanoic acid (C8), an 8-carbon linear monocarboxylic acid, is a particularly valuable MCFA because of its high market value and relative ease of recovery. Further, when C8 is reduced to its corresponding alkane, it can substitute for octane in liquid transportation fuels. Recent research has demonstrated the feasibility of using mixed culture fermentations to transform renewable resources into MCFAs.^5-7^ While C8 is produced by some self-assembled bioreactor microbial communities, hexanoic acid (C6) is often the primary MCFA, along with two short-chain fatty acids (SCFAs), acetic (C2) and butyric (C4) acids.^5-10^

The microbial metabolic networks involved in converting complex organic substrates to MCFAs by mixed culture fermentations are not fully elucidated. Consequently, there is a lack of knowledge on how to obtain a carboxylate platform community that produces C8 as the main MCFA, or how to minimize the accumulation of SCFAs. Initial models of mixed culture fermentations to produce products other than methane have focused on simulating metabolic networks for production of C2, C4, propionic acid, ethanol, lactic acid and H_2_ from glucose.^11^ Subsequent models took into account that multiple electron carriers (e.g., ferredoxin, NADH) play different roles in these metabolic networks, enabling the simulation of energy-conserving electron-bifurcating reactions during production of C4 by reverse β-oxidation.^12^ While a recent model considered the role of homoacetogens as a H_2_-consuming member of mixed culture fermentation, it was limited to producing C4 as the major product of reverse β-oxidation and did not include MCFA production.^13^ Additionally, recent metabolic models assessed the potential for producing C2, C3, C4 and valerate (C5) from amino acids.^14^ While all of these models have enabled predictions of mixed-culture fermentations from a single organic substrate, they lack the ability to simulate production of MCFAs from complex feedstocks.

In this study, we constructed a unicellular model (iFermCell215) and a guild-based microbial community model (iFermGuilds789) that offer expanded substrates and metabolic pathways compared to existing models. We used iFermCell215 and iFermGuilds789 to analyze pathways that lead to MCFA production. Simulations were performed with single organic substrates, including sugars, glycerol, lactate, and acetate, with ethanol and acetate or lactate and acetate as co-substrates, and with complex organic mixtures. The iFermGuilds789 was also used to evaluate predictions of observed bioreactor behavior and of the flow of carbon and electrons through different guilds within the mixed microbial community. In total, results of individual model simulations generated new hypotheses on how to increase MCFA production by mixed microbial communities.

## MATERIALS AND METHODS

Metabolic networks of mixed culture fermentation that contain reactions that occur during anaerobic metabolism of a previously described lignocellulosic biorefinery residue^15^ were assembled. The first network, iFermCell215, contains 215 reactions and 154 metabolites, models mixed culture fermentation as a single unit, and expands the metabolic pathways and substrate range of previously published mixed culture fermentation networks.^11-13^ iFermCell215 describes metabolism of exogenous substrates including glucans, glucose, xylans, xylose, and glycerol, along with lactate, ethanol, acetate, CO_2_ and H_2_, which are potential intermediates in this mixed culture fermentation (**Fig. 1**).

**Figure 1.**
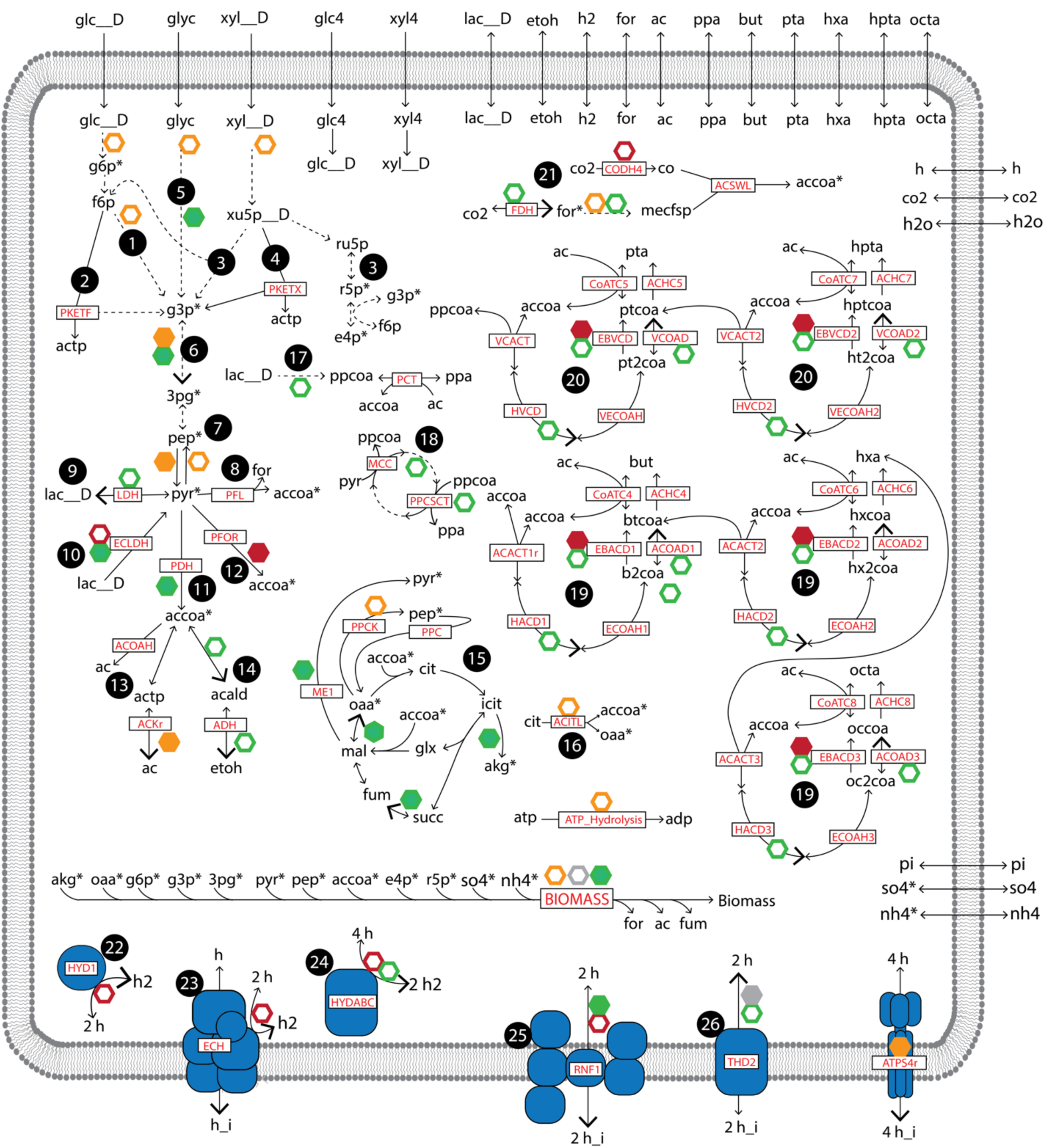
Summary of metabolic networks in iFermCell215. Metabolites with an asterisk are required for biomass production. Open hexagons indicate consumption of ATP (orange), NADH (green), NADPH (grey), and reduced ferredoxin (red). Closed hexagons indicate production of these compounds and consumption of ADP (orange), NAD+ (green), NADP+ (grey), and oxidized ferredoxin (red). For reversible reactions, larger arrowheads indicate the reaction directionality corresponding with consumption and/ or production of ATP, NADH, NADPH and reduced ferredoxin.. Dashed lines indicate multiple reactions. Key reactions are indicated according to the abbreviation used in iFermCell215 (**Supplementary File 3**). Pathways are identified by number: (1) upper glycolysis; (2) phosphoketolase metabolism of fructose; (3) non-oxidative pentose phosphate pathway; (4) phosphoketolase metabolism of xylulose; (5) glycerol utilization; (6) lower glycolysis; (7) gluconeogenesis; (8) pyruvate formate lyase; (9) lactate metabolism; (10) pyruvate flavodoxin oxidoreductase; (11) pyruvate dehydrogenase; (12) acetate metabolism; (13) ethanol metabolism; (14) incomplete citric acid cycle with glyoxylate shunt; (15) anaplerotic metabolism of phosphoenolpyruvate, pyruvate, and oxaloacetate; (16) ATP citrate lyase; (17) propionate production via the acryloyl-CoA pathway; (18) propionate production via the methylmalonyl-CoA pathway; (19) even-chain reverse β-oxidation; (20) odd-chain reverse β-oxidation; (21) homoacetogenesis via the Wood Ljungdahl pathway; (22) ferredoxin-dependent hydrogenase; (23) proton-translocating hydrogenase; (24) electron confurcating hydrogenase; (25) proton-translocating ferredoxin: NAD^+^ oxidoreductase; (26) NAD(P) transhydrogenase. For clarity, not all reactions or metabolites are shown. More information on the reactions and metabolites contained in iFermCell215 is provided in **Supplementary File 3**.

The second network, iFermGuilds789, represents activities of different guilds (i.e., groups of organisms with similar functions) within the microbial community. The iFermGuilds789 network describes the same reactions and metabolites as in the iFermCell215 model but these are separated into six functional guilds with each guild representing a subset of compartmentalized reactions (**Fig. 2**). iFermGuilds789 also includes additional transport reactions that simulate metabolite exchange among guilds. Combining the six guilds, the iFermGuilds789 network includes 789 reactions and 579 metabolites. The guilds in the iFermGuilds789 model represent previously described organisms that perform fatty acyl chain elongation from sugars (SEOs), sugar fermenting organisms that produce lactate, acetate, and ethanol as fermentation products (SFO), hydrogen-producing sugar fermenters (HSFs), organisms that perform fatty acyl chain elongation from lactate (LEOs) or ethanol (EEOs), and homoacetogenic organisms (HAOs). Examples of organisms falling within each functional guild are described in **Supplementary Text 1**.

**Figure 2.**
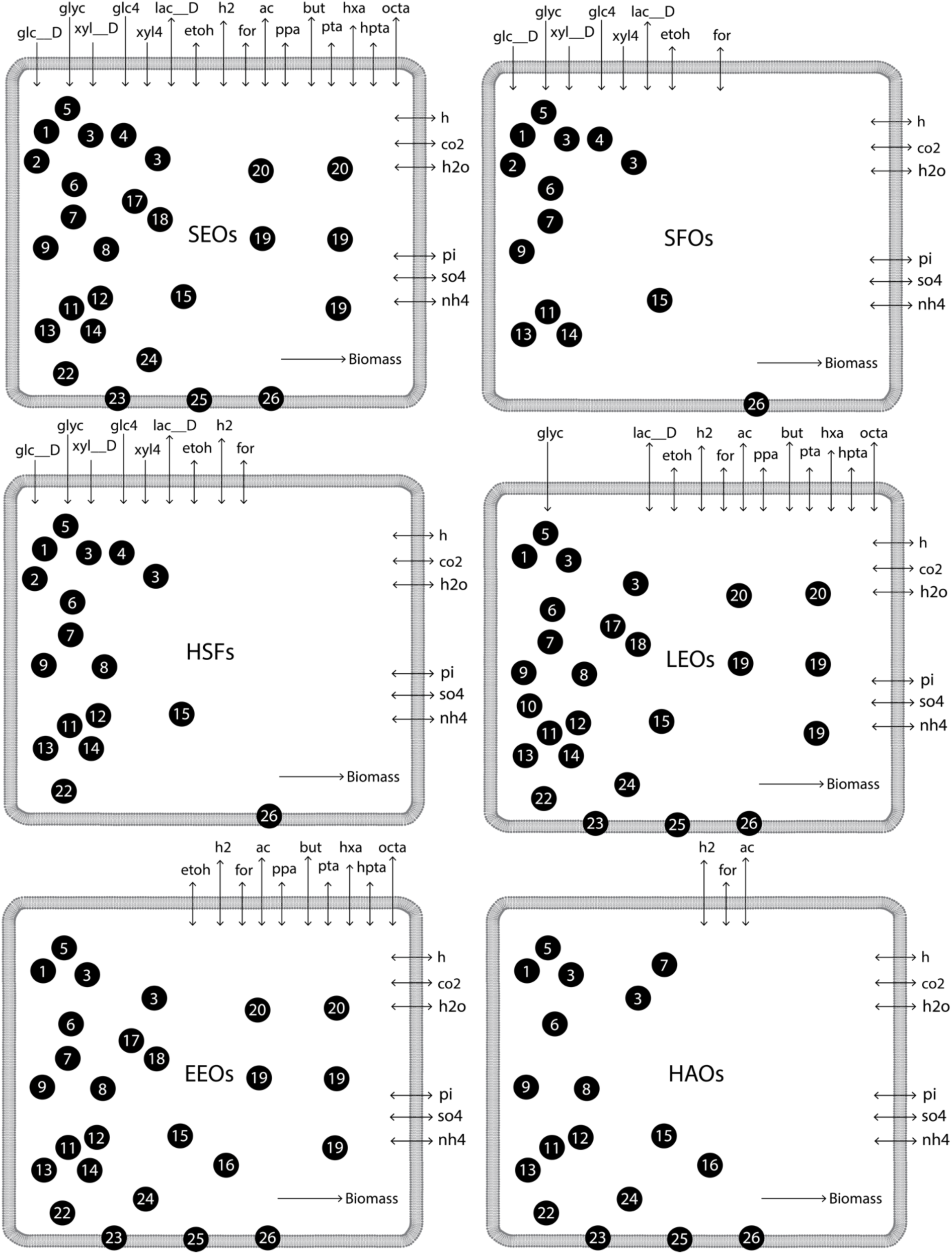
Summary of metabolic networks for six functional guilds contained in iFermGuilds789. Refer to **Fig. 1** for details on pathways included in the iFermGuilds789 model. More information on the reactions and metabolites contained in the iFermGuilds789 model is provided in **Supplementary File 4**.

When constructing each metabolic network, reactions were added to capture the exchange of substrates and products with the extracellular space.^16^ In addition to intracellular (denoted as “_c” in the metabolic models) and extracellular (“_e”) compartments, a compartment for ion-motive force generation was included (denoted as “_i” in the metabolic models). Where possible, standard metabolite and reaction abbreviations from the Biochemical, Genetic, and Genomic (BiGG) knowledgebase were used.^17^ All reactions were balanced for mass and charge assuming an intracellular pH of 7.0 with the dominant ionic form of all metabolites obtained from the BiGG database.^17^

Since many members of this mixed microbial community are Firmicutes, simplified biomass equations for both models were created based on the biomass composition of the well-studied gram-positive bacterium *Bacillus subtilis*.^*18-20*^ We simplified all of the biosynthetic pathways for production of biomass components to capture 16 precursor compounds produced by the models. The precursor compounds included α-ketoglutarate, oxaloacetate, glucose-6-phosphate, glucose-3-phosphate, 3-phosphoglycerate, pyruvate, phosphoenolpyruvate, acetyl-CoA, erythrose-4-phosphate, ribulose-5-phosphate, ATP, NADPH, NAD+, ammonium, sulfate, and H_2_O. Additional information and calculations for the biomass equation are provided in **Supplementary Text 2, Supplementary Data File 1, Figures S1-S6, and Table S1**. Both metabolic models (iFermCell215 and iFermGuilds789) and instructions for their use are available at https://github.com/mscarbor/Mixed-Culture-Fermentation-Models.

The production capabilities of both metabolic networks were simulated using constraint-based methods.^21^ These methods predict the steady-state flux distribution through a metabolic network that is subject to physiochemical constraints. In this work, we used flux sampling^22^ and parsimonious flux balance analysis (pFBA).^23^ Flux sampling was performed using artificial centering hit and run (ACHR)^22^ using 101 simulations and randomly generated seeds. A solution with pFBA provides a flux distribution by minimizing the sum of all fluxes predicted by the model.^23^ When modeling bioreactors with the presence of solute gradients across the cell membrane, we considered the ATP required for transport as described in **Supplementary Text 3**. The objective function for all modeling scenarios was to maximize biomass production. When simulating the iFermGuilds789 network for a steady-state condition, we required all individual guilds to achieve the same growth rate (hr^-1^). All simulations were performed in Python 3.7 using the cobrapy package (v 0.18.1).^24^

Throughout this work, we present results in terms of electron equivalents (eeq), which report the number of electrons potentially transferred to an electron acceptor if the compound is completely oxidized (using factors provided in **Table S2** to convert from molar concentrations to eeq concentrations). For each scenario, we calculated the overall reaction thermodynamics using the predicted stoichiometry of the model and the standard free energy of formation (ΔG_f_^0^) of metabolites (**Table S2**) obtained from KBase.^25^ Unless otherwise noted, we ensured that at least 50 kJ was available per ATP produced under standard conditions at neutral pH.^26, 27^

## RESULTS

### Simulation of mixed culture fermentations with iFermCell215

By constraining the iFermCell215 model to produce defined products, we initially tested its predictive ability with well-established fermentation conditions (See **Supplementary Text 4** and **Table S3**). We then investigated predictions derived from an iFermCell215 model under conditions where single substrates were consumed within this unit. The iFermCell215 model depicts a simplified representation of fermentation in a mixed microbial community where interspecies transfer of metabolites is assumed to have no energetic penalty, and is similar to previously published models of mixed culture fermentations.^11-13^ For this analysis, we required that all possible solutions released ≥ 50 kJ per mol ATP produced.^26^

Using flux sampling, iFermCell215 provided 101 solutions that maximized biomass production depending on the substrates provided to the simulation (**Fig. 3**). For instance, when we modeled the utilization of glucose or xylose as the sole substrate, iFermCell215 predicted multiple solutions that allowed for maximum biomass production (40% of eeq are incorporated into biomass) with H_2_, C2, C4, C6, and C8 as predicted products (**Fig. 3A**). None of the solutions in this flux sampling analysis predicted ethanol or lactate as significant products. The predicted biomass yield (eeq incorporated into biomass per eeq consumed) decreased with glycerol (37%), lactate (20%), and ethanol (5.9%) as sole substrates. Consumption of ethanol as a sole substrate was predicted to produce C8 and H_2_ as the most energetically favorable solution (**Fig. 3D**), whereas consumption of lactate as a sole substrate produced optimal solutions through heterofermentation (**Fig. 3C**) and consumption of H_2_ and CO_2_ was predicted to produce C2 for optimal biomass growth, which is consistent with the activity of homoacetogens (**Fig 3E**). These results suggest that sugars, glycerol, and lactate can produce significant amounts of multiple fermentation products, including C2, C4, C6, and C8, while supporting optimal biomass production, and that production of ethanol or lactate, the two common fermentation products described as intermediates in MCFA production,^28^ are not as energetically favorable as producing other fermentation products. It further suggests that consumption of ethanol as a sole substrate should favor production of C8, and therefore, that ethanol may be a better intermediate product for optimizing C8 production than lactate. However, since the fermentation of complex organic substrates by mixed communities often results in accumulation of multiple fermentation products, we also simulated conditions in which ethanol and acetate or lactate and acetate were co-substrates for MCFA production.

**Figure 3.**
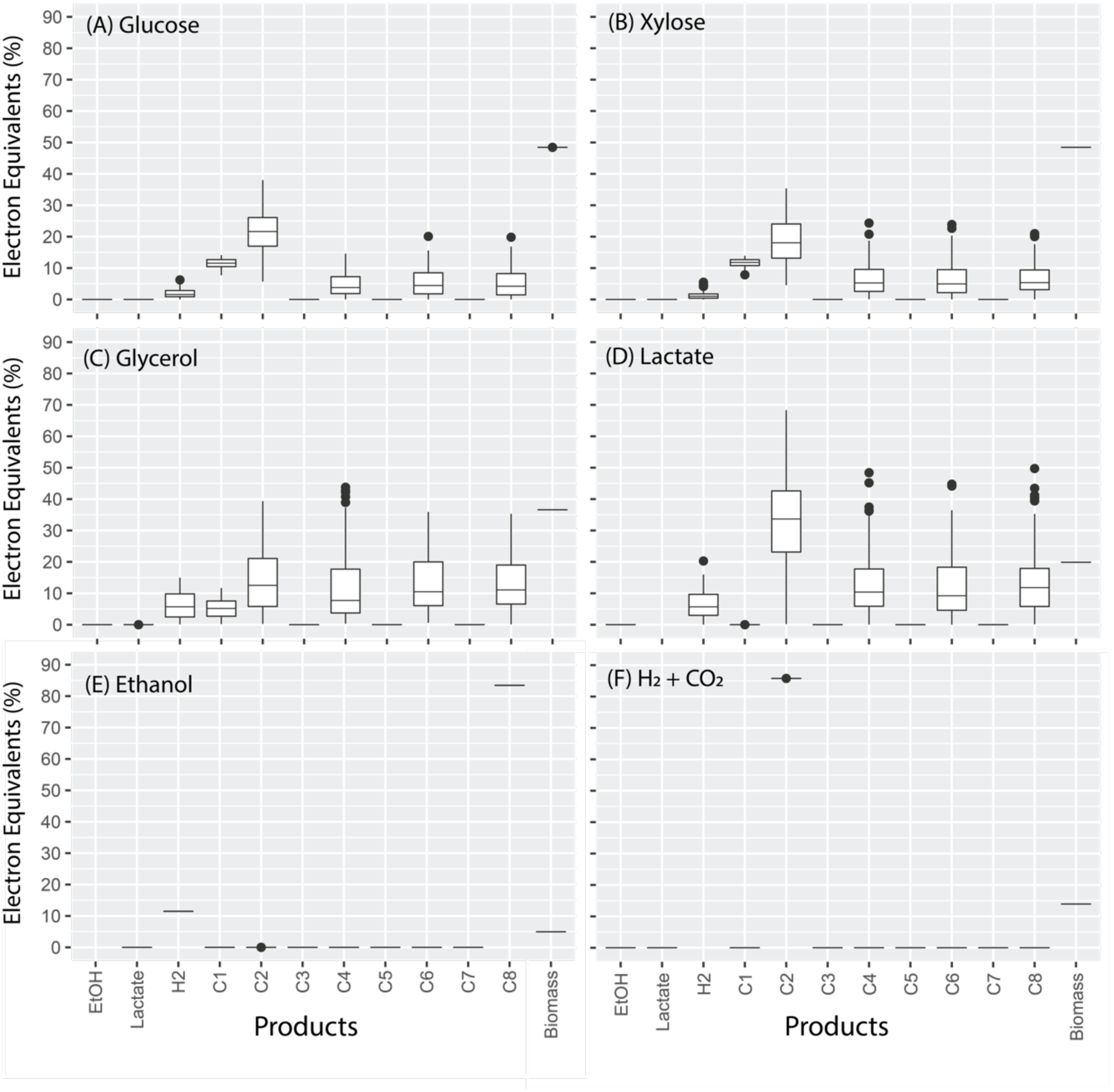
Flux sampling results for iFermCell215 with objective function to optimize biomass production when fed substrates indicated within the plots: (A) glucose, (B) xylose, (C) glycerol, (D) lactate, (E) ethanol, (F) H_2_ and CO_2_ (with H_2_ as the limiting substrate). Box plots show the results of 101 simulations. Median values are indicated by a horizontal line within the box, and the top and bottom of the box indicate the 25^th^ and 75^th^ percentile, respectively. Whiskers indicate values within 1.5 times the interquartile range and closed circles indicate values outside of this range.

### Simulation of ethanol and acetate utilization with iFermCell215

Ethanol is a substrate that has often been used for chain elongation,^29^ and early work on MCFA production was performed using the ethanol- and acetate-consuming *Clositridium kluyveri*.^30, 31^ During initial work with *C. kluyveri*, it was proposed that C4 could be produced from two moles of ethanol, without using acetate as an electron acceptor.^31^ Recent work has confirmed that *C. kluyveri* produces MCFAs from ethanol as a sole substrate in batch experiments,^32^ in agreement with the iFermCell215 prediction (**Fig. 3E**). Production of MCFAs from C2 as a sole substrate has not been observed and is not predicted by iFermCell215. Past studies of ethanol elongation have investigated optimal ratios of ethanol-to-acetate for MCFA production.^29, 33, 34^ Here, we use iFermCell215 to assess impacts of different molar ratios of acetate-to-ethanol (C2:EtOH) on predicted product formation.

When supplied with both ethanol and C2, the iFermCell215 predicted maximum growth rate of 0.11 hr^-1^ is more than double the predicted growth rate with ethanol as the sole substrate (**Fig. 4A**), suggesting an ecological advantage for microorganisms that can co-utilize ethanol and acetate. This growth rate can be achieved at C2:EtOH ratios from 0.56 to 1.28 mol:mol. At ratios of 0.56 mol:mol and lower, C8 and H_2_ are the only major predicted products (**Fig. 4**, left panels). At ratios between 0.56 and 1.20, a mixture of C8, C6, and C4 are predicted by iFermCell215, and at ratios of 1.28 and above, C4 is predicted as the sole product. These model predictions are consistent with previous experimental work that shows that low ratios of C2:EtOH improve specificity of C8 production.^33^ The overall production rate of MCFAs, however, has been shown to improve at higher ratios of C2:EtOH.^33^ The highest overall MCFA production rate was achieved by Spiritio et. al at a ratio of 0.53 mol:mol,^33^ which is very close to the model predicted ratio of 0.56 that optimizes biomass growth and C8 production. The production of H_2_ at low ratios of C2:EtOH raises the question as to the impact of H_2_ partial pressure on the thermodynamic feasibility of the model results. We used iFermCell215 to calculate the maximum H_2_ partial pressure for all 101 scenarios and found that the lowest allowable H_2_ partial pressure for growth is greater than 10^6^ atm (**Fig S7**), indicating that the proposed transformations are thermodynamically favorable under all bioreactor H_2_ partial pressure conditions.

**Figure 4.**
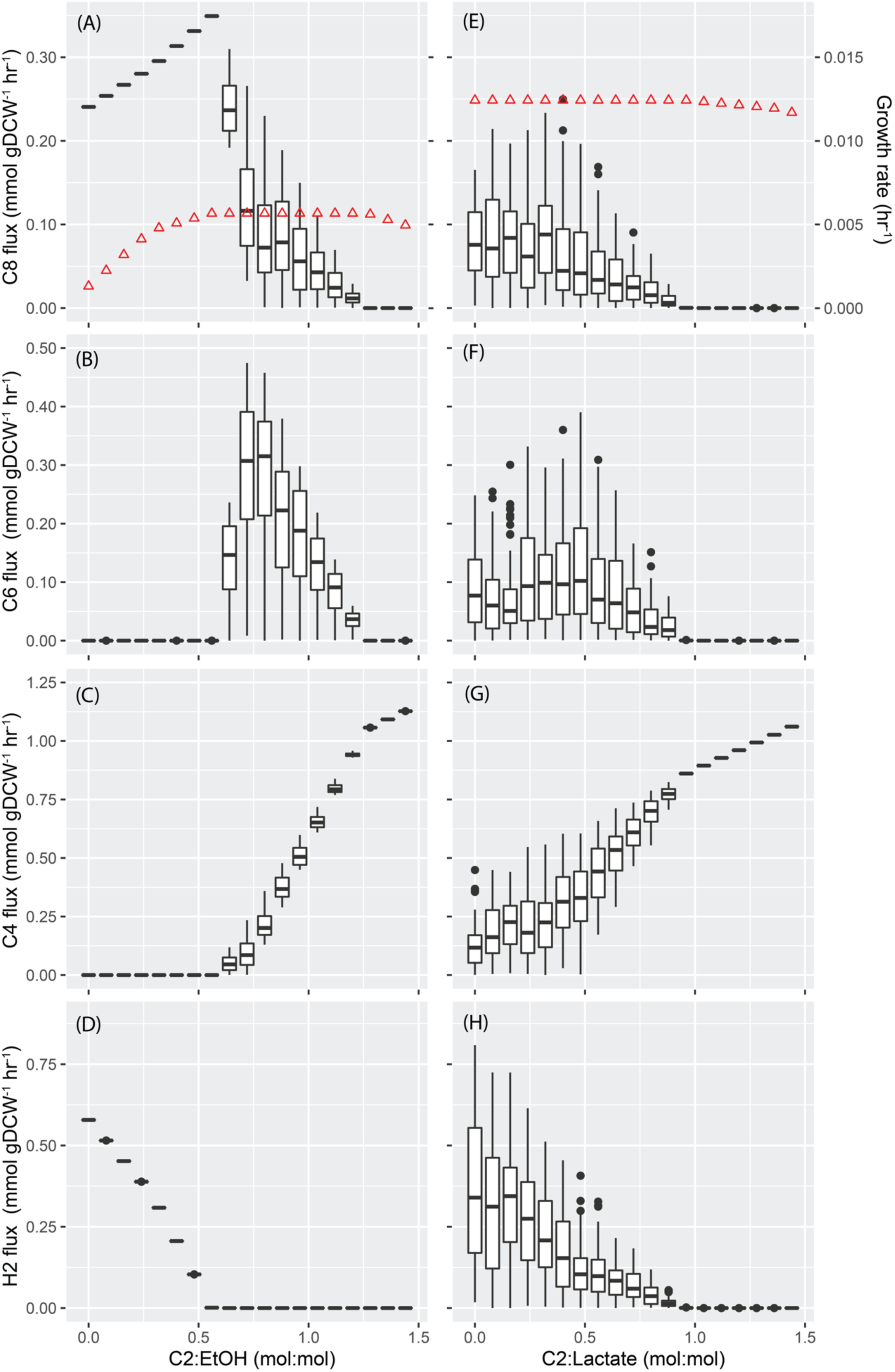
Flux sampling results for iFermCell215 when consuming ethanol and acetate (A-D) or lactate and acetate (E-H) at different molar ratios of acetate-to-ethanol (C2:EtOH) or acetate-to-lactate (C2:Lactate). Results are shown for fluxes of octanoate (C8), hexanoate (C4), butyrate (C4) and H_2_. These results are shown as box plots, as described previously (Figure 3), representing the results of 101 simulations for each modeled ratio. The predicted maximum growth rate (hr^-1^) is shown for ethanol (A) and lactate (E) as red triangles plotted on the secondary y-axis.

In total, iFermCell215 predicts that the C2:EtOH ratio could either be used to improve production of C8 as a desired product, or could be an indication that the environmental conditions in a bioreactor would lead to a mixture of fermentation products. At low C2:EtOH ratios (<0.56), iFermCell215 predicts that C8 should be the abundant product. At medium C2:EtOH ratios (0.56-1.20) iFermCell215 predicts a mixture of C4, C6, and C8 would be produced, while at high ratios (>1.20) of these substrates, iFermCell215 predicts C4 would be the predominant product. Therefore, in a microbial community fermenting a mixture of substrates, iFermCell215 predicts that different ratios of ethanol and C2 will impact the composition and abundance of products. Additionally, iFermCell215 predicts that, if feeding a reactor with ethanol and C2, attention should be paid to creating a feeding regime that optimizes production of the target product.

### Simulation of lactate and acetate utilization with iFermCell215

In addition to ethanol, lactate is often used as a substrate for microbial chain elongation. The well-studied *Megasphaera elsdenii* has been shown to produce MCFAs from lactate,^35^ and recent chain elongation studies with microbial communities have explored lactate as both a substrate^9, 36^ and a proposed key intermediate that supports chain elongation from carbohydrate fermentation.^15, 37^ While ethanol and lactate are both potential fermentation products and substrates for chain elongation, iFermCell215 predicts that the pathways for utilization of these substrates have key differences that impact the terminal fermentation products, as shown in the flux sampling analyses (**Fig. 3D-E**), and may impact the products when C2 is a co-substrate. Therefore, similar to the analysis shown above for C2:EtOH ratios, we used iFermCell215 to assess impacts of different C2-to-lactate ratios.

In the case of lactate, the predicted maximum growth rate does not change when C2 is a co-substrate (**Fig. 4E**). This suggests that, unlike ethanol and C2 co-utilization, there is no apparent energetic benefit (reflected in growth rate) for a microorganism to co-utilize lactate and C2. Low ratios of C2:lactate are predicted by iFermCell215 to produce a variety of end products, including C8, C6, C4, and H_2_ (**Fig. 4**, right panels). As the C2:lactate ratio increases, the product distribution is predicted by iFermCell215 to change, with a decrease in MCFA and H_2_ production accompanied by an increase in C4 production, and at ratios of 0.96 and above, C4 is predicted as the sole product. Like ethanol and C2 co-utilization, lactate and C2 co-utilization is not predicted by iFermCell215 to be impacted by bioreactor H_2_ partial pressure (**Fig S7**). These results suggest that when lactate is a key intermediate in an MCFA-producing community, controlling the relative amounts of C2 and lactate may not be an effective strategy for increasing C6 and C8 production. However, like with ethanol, an excess amount of acetate is predicted by iFermCell215 to increase C4 production.

The predicted differences in product formation when acetate is co-metabolized with ethanol or lactate suggests important differences in the metabolism of these substrates. Therefore, to understand these differences, we also used iFermCell215 to assess the predicted metabolic pathways of ethanol and lactate utilization.

### Predicted metabolic pathways for ethanol and lactate utilization

The flux sampling analysis obtained with iFermCell215 consuming ethanol as a sole substrate converged to a solution in which C8 was the only significant fermentation product (**Fig. 3E**). The flux of electron equivalents in this simulation predicted that reactions catalyzed by ferredoxin hydrogenase (Hyd1), proton-translocating ferredoxin:NAD+ oxidoreductase (aka, *Rhodobacter* nitrogen fixation complex; RNF), a coenzyme A transferase (CoAT), and acetate kinase (AckR) were key enzymes used in the network (**Fig 5A**). The genome of the well-studied ethanol chain-elongating organism, *C. kluyveri*, is predicted to contain genes that encode the proteins for catalyzing all of these reactions.^38^ Further, Hyd1 is the only hydrogen-producing gene annotated in the genome of *C. kluyveri*.

**Figure 5.**
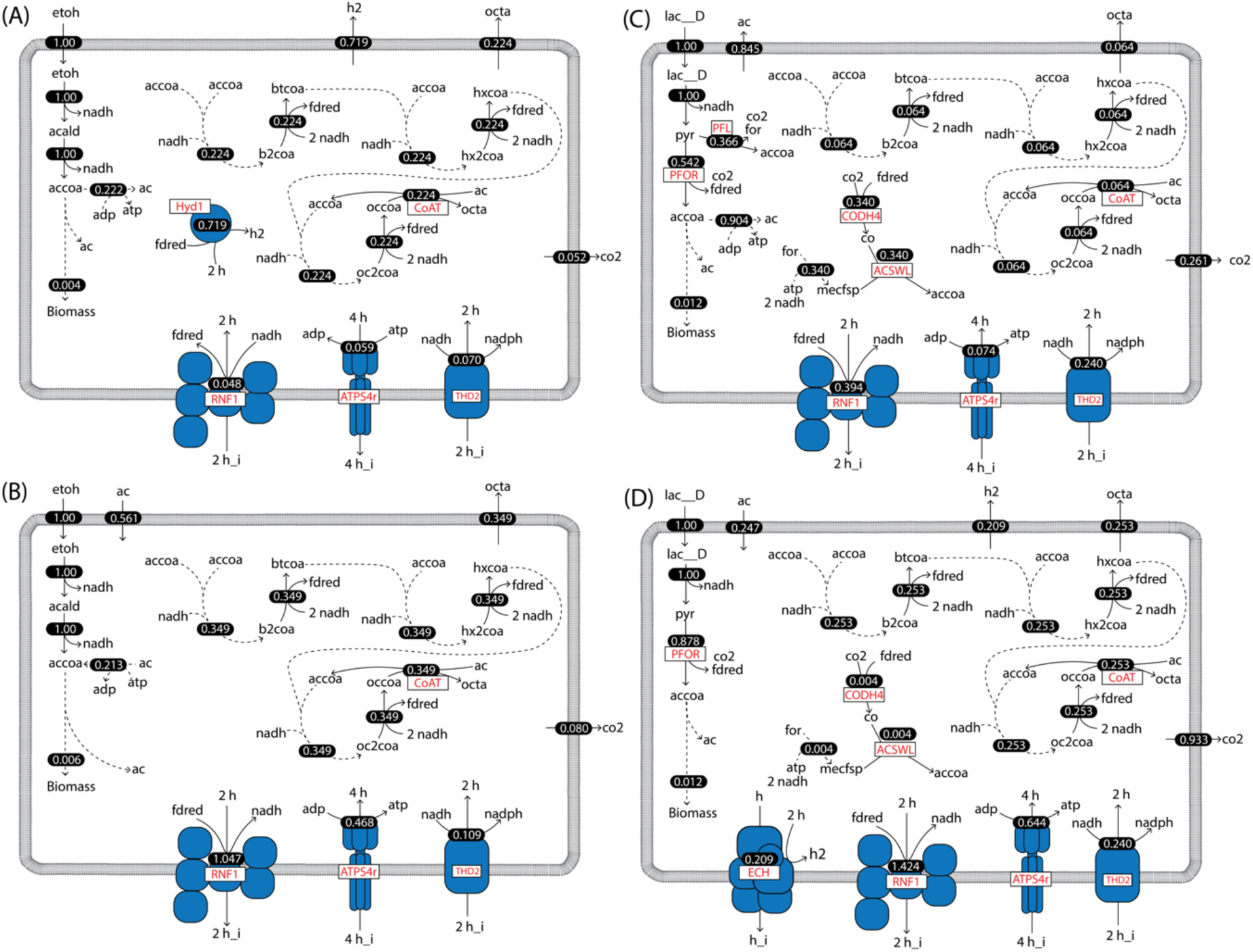
Predicted pathways of ethanol utilization by iFermCell215 using pFBA when (A) ethanol is the sole substrate; (B) ethanol and acetate are co-substrates; (C) lactate is a sole substrate; and (D) lactate and acetate are co-substrates. Flux values (shown as white text) have units of mmol gDCW^-1^ hr^-1^. For both scenarios, the model was constrained to uptake 1 mmol gDCW^-1^ hr^-1^ of ethanol or 1 mmol gDCW^-1^ hr^-1^ of lactate. For scenarios with acetate co-utilization, acetate was provided in excess. Key reactions indicated by red text include a ferredoxin hydrogenase (Hyd1), a CoA transferase (CoAT), a proton-translocating ferredoxin: NAD+ oxidoreductase (RNF1), ATP synthase (ATPS4r), NADH:NADP transhydrogenase (THD2), and energy-conserving hydrogenase (ECH). Metabolite abbreviations are and are defined in **Supplementary Data File 2**.

The flux through the above enzymes predicts extensive use of energy conserving mechanisms that rely on the generation of an ion-motive force (IMF). When ethanol is the sole substrate (**Fig. 5A**), iFermCell215 predicted that the IMF is generated by the ATP synthase while hydrolyzing ATP. The model predicts that the RNF enzyme uses the IMF to generate reduced ferredoxin and that ATP is provided by acetate kinase. The reverse β−oxidation cycle uses an electron bifurcating acyl-CoA dehydrogenase that also generates reduced ferredoxin, and redox balance is maintained by Hyd1 that uses the reduced ferredoxin to generate H_2_ **(Fig. 5A)**. When C2 is available in addition to ethanol, iFermCell215 predicts that no net H_2_ production is required for redox balance **(Fig. 5B)**. Instead, iFermCell215 predicts that the activities of several enzymes are reversed, with the RNF complex generating IMF, ATP synthase producing ATP, and acetate kinase consuming ATP to generate acetyl-CoA. Thus, the same set of reactions is predicted to be important by iFermCell215 when using ethanol as a sole substrate or ethanol and C2 as co-substrates, but the predicted direction of key reactions change (**Fig 5A-B**). These predictions suggest that organisms that elongate ethanol may readily transition between utilizing ethanol as a sole substrate and co-utilizing ethanol and C2 since the same enzymes are predicted to be used, and the co-utilization of C2 and ethanol improves growth rates (**Fig 4A**).

Unlike ethanol utilization, lactate conversion to acetyl-CoA is predicted by iFermCell215 to produce CO_2_, formate, and reduced ferredoxin (**Fig. 5C-D**). The Wood-Ljungdahl pathway is also predicted by iFermCell215 to use this CO_2_, formate and reduced ferredoxin to produce C2 (**Fig. 5C-D**). The predicted ability to generate reduced ferredoxin during lactate consumption (compared to only NADH from ethanol consumption) enables iFermCell215 to utilize a much wider range of pathways for generating energy and maintaining redox balance. The reverse β−oxidation cycle also uses an electron bifurcating acyl-CoA dehydrogenase under these conditions, generating reduced ferredoxin, with RNF also predicted to be active in redox balancing. These predictions of iFermCell215 explain the wide range of products that are possible from utilization of lactate as sole substrate and co-utilization of C2 and lactate. Further, this suggests that limiting the available metabolic pathways can impact the predicted products from lactate. Indeed, removing specific reactions in iFermCell215 impacts the predicted product spectrum (**Supplementary Text 5, Fig S7**). This demonstrates that strategies to improve MCFA production may depend on the available set of microbial transformations and may differ between bioreactor microbial communities.

### Simulating community behavior with iFermCell215 and iFermGuilds789

The iFermCell215 model predicted that multiple substrates could lead to C8 production (**Fig. 3**), but that biomass could be maximized by co-production of other fermentation products. We wanted to assess if this single-unit model could simulate the observed product yields of a mixed microbial community fed lignocellulosic biorefinery residues, a complex organic substrate.^15^ The previously described bioreactor was fed lignocellulosic biorefinery residues for 96 days, at which time its microbial community was evaluated using metagenomic and metatranscriptomic analyses along with end-product sampling. To represent organic substrate metabolism, the substrate uptake fluxes were set according to the observed consumption of compounds in the feedstock (**Table S6**). We constrained iFermCell215 by setting the growth rate of the model to 0.0069 hr^-1^, corresponding to the bioreactor retention time of 6 d and we again constrained the model only by requiring a minimum of 50 kJ of free energy release per mol of ATP produced. Flux sampling results from this scenario predicted a range of potential products (**Fig. 6A)**, including ethanol, C2, H_2_, even chain carboxylic acids, and odd-chain carboxylic acids. Overall, the amount of MCFAs predicted by iFermCell215 exceeded observed values, exposing a limitation of the single unit model.

**Figure 6.**
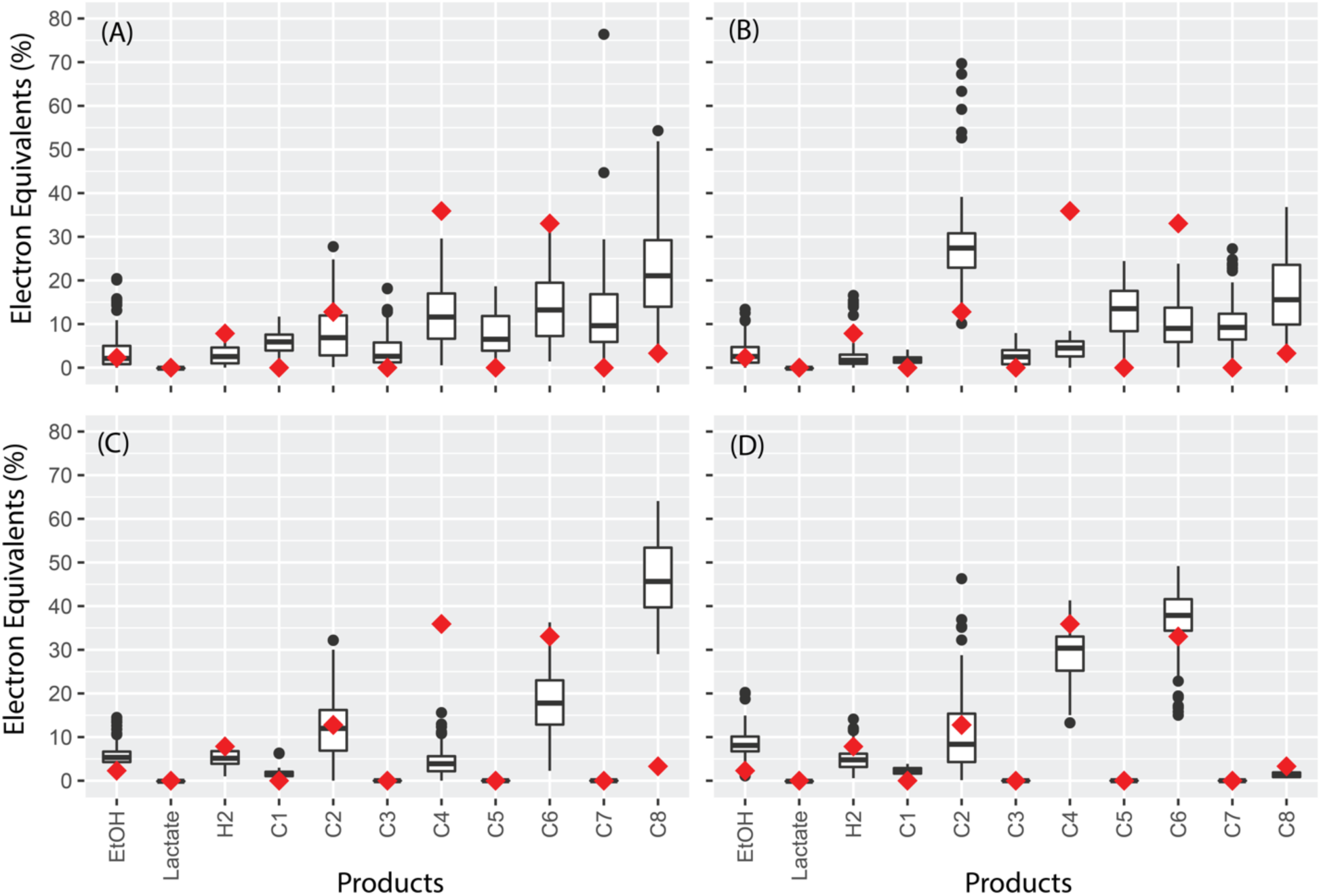
Predicted and observed product formation for a mixed culture fermentation bioreactor fed lignocellulosic biorefinery residues. (A) predictions with iFermCell215; (B) predictions with iFermGuilds789; (C) predictions with iFermGuilds789 with reactions knocked-out based on metagenomic analyses of the microbial community; (D) predictions with iFermGuilds789 with additional constraints for maximum fluxes through acyl-CoA and CoAT reactions for C6 and C8 production in SEOs and LEOs

In an attempt to better represent the compartmentalized nature of activities within mixed microbial communities we developed the functional guild model iFermGuilds789. In iFermGuilds789, the metabolic activities of a mixed microbial community are partitioned among a number of functional guilds. This biological division of labor negates the free exchange of reducing equivalents (e.g., NADH, ferredoxin) and of many pathway intermediates (e.g., acetyl-CoA) between community members, ultimately constraining how individual organisms maintain redox balance, which can potentially impact ATP and product yields. For instance, the single unit model (iFermCell215) cannot predict the production of lactate via a NADH-dependent dehydrogenase and re-consumption of lactate via an electron-confurcating dehydrogenase, activities that may happen in a mixed microbial community that includes lactate producers and lactate consumers. The iFermGuilds789 model contains fermentation pathways in six guilds (**Fig. 2**), with each guild including predictions on transport reactions to simulate uptake and release of intermediate metabolites by the most abundant members of the community as predicted from previous metagenomics analyses.^15^

To simulate the activities in the MCFA-producing bioreactor we used a version of iFermGuilds789 with only four guilds, which represented the most abundant community members in the bioreactor as determined by metagenome sequencing.^15^ These guilds included the sugar elongating Cand. *Weimeria bifida* (SEOs), lactate-elongating Cand. *Pseudoramibacter fermentans* (LEOs),^39^ five sugar fermenting *Lactobacillus* species (SFOs), and three hydrogenic sugar fermenting *Coriobacteriaceae* species (HSFs).^*15*^ We constrained the abundance of each guild in iFermGuilds789 according to the relative abundance of DNA for each guild in the bioreactor. Guilds were assumed to have all pathways associated with their guild (**Fig. 2**), and we required each guild to have the same growth rate of 0.0069 hr^-1^. The results of this iFermGuilds789 simulation (**Fig. 6B**) predicted a narrower range of products than iFermCell215 (**Fig. 6A**), with even- and odd-chain fatty acid products predicted. With these parameters, the iFermGuilds789 model overpredicted the production of C2 and C8 while underpredicting C4 and C6 production.

In an additional iFermGuilds789 simulation, we restricted pathways in guild members based on previous metatranscriptomic analyses.^*15*^ SEOs were constrained to use glucose, xylose and glycerol; SFOs were restricted to consuming glucose, xylose, xylans, glucans, and glycerol; HSFs were constrained to only consume glucans, glucose and glycerol; and LEOs were restricted to consuming only lactate and glycerol (**Table S7**). Further, SEOs and LEOs were constrained to not produce odd-chain volatile fatty acids (propionate, pentanoate, and heptanoate). LEOs were further constrained to not use acetate kinase, which is absent from the genome of Cand. *Pseudoramibacter fermentans*.^39^ This iFermGuilds789 simulation predicted ranges of ethanol, lactate, C2, and C6 that contained the experimentally observed values (**Fig 6C**). With these parameters, C8 and H_2_ production were overpredicted by iFermGuilds789, while C4 production was underpredicted.

One potential explanation for the differences in the various iFermGuilds789 predictions may be the specificity of chain elongating enzymes for producing products of defined chain lengths. This could include any of the enzymes catalyzing the four steps of reverse β-oxidation itself or the terminal enzymes that cleave the final products of reverse β-oxidation from CoA. We tested this by constraining the flux through CoAT and acyl-CoA hydrolase reactions for both SEOs and LEOs according to carbon-chain length. When constraining both CoAT and acyl-CoA hydrolase in both SEOs and LEOs to 0.02 mmol gDCW^-1^ hr^-1^ for C8 and 1 mmol gDCW^-1^ hr^-1^ for C6, iFermGuilds789 modeling predictions aligned much better with community performance (**Fig. 6D**). This suggests that reverse β-oxidation and/or terminal enzymes in this pathway have higher affinity for C4 and C6 than for C8 in this specific microbial community.

Taken together, initial predictions from iFermCell215 (**Fig. 6A**) and iFermGuilds789 (**Fig. 6B**) did not align with observed bioreactor performance. Predictions of iFermGuilds789 improved by incorporating knowledge of specific reactions within each guild (**Fig. 6C**). Further, the best alignment between observations and model predictions resulted from constraining the fluxes related to individual products with iFermGuilds789 (**Fig. 6D**). This suggests that accurate predictions of MCFA production may require incorporation of enzyme affinity and additional characterization of enzymes from chain-elongating species to improve modeling predictions.

### Assessing interspecies metabolite flux using iFermGuilds789

Our previous analyses predicted that microbes within the MCFA-producing microbial community perform different functions, compete for resources, and participate in interspecies transfer of metabolites that allows for conversion of lignocellulosic biorefinery residues to SCFAs and MCFAs.^15^ We used iFermGuilds789 to predict the function of each guild in the transformation of substrates as well as the production and transfer of intermediate metabolites between microbes. In this case, unlike previous simulations with iFermGuilds789, we did not constrain the growth rate (**Fig. 6B-D**) or fluxes for MCFA production (**Fig. 6D**). Instead, we constrained iFermGuilds789 to accumulate extracellular products at the rates observed in the MCFA-producing bioreactor (**Table S6**), constrained the reactions within each guild as described above, and predicted fluxes using pFBA, which calculates a single flux distribution when the model is constrained to the observed production rates. Under these conditions, iFermGuilds789 predicted a growth rate of 0.0051 hr^-1^, which is 74% of the expected growth rate (0.0069 hr^-1^). This growth rate corresponds to a retention time of 8 days (compared to the actual retention time of 6 days) and these differences may reflect difficulty in estimating the amount of metabolically active cells in the reactor or absence of unknown metabolic pathways that may increase the predicted growth rate.

By constraining the substrates and products according to observed community behavior, iFermGuilds789 predicted that SFOs and HSFs consumed xylans and glucans, while SEOs consumed xylose, glucose, and 99% of the glycerol (**Fig 7**). SFOs were predicted to produce ethanol, lactate and C2, while HSFs were predicted to produce a small amount of the lactate and C2. SEOs are also predicted to produce lactate and C2. In total, 22% of the eeq in the feedstock are predicted by this instance of iFermGuilds789 to be converted to lactate and 27% are predicted to be converted to C2. All of the lactate and 52% of the C2 is predicted to be consumed by LEOs. LEOs are predicted to produce all of the C4. SEOs are predicted to produce all of the C6 and C8 along with all of the H_2_. SEOs are further predicted to have a flux for C6 production of 0.46 mmol gDCW^-1^ hr^-1^ and C8 production of 0.033 mmol gDCW^-1^ hr^-1^ (**Supplementary Data File 5**), both of which are lower than the maximum flux constraints applied previously (**Fig. 6D)**.

**Figure 7.**
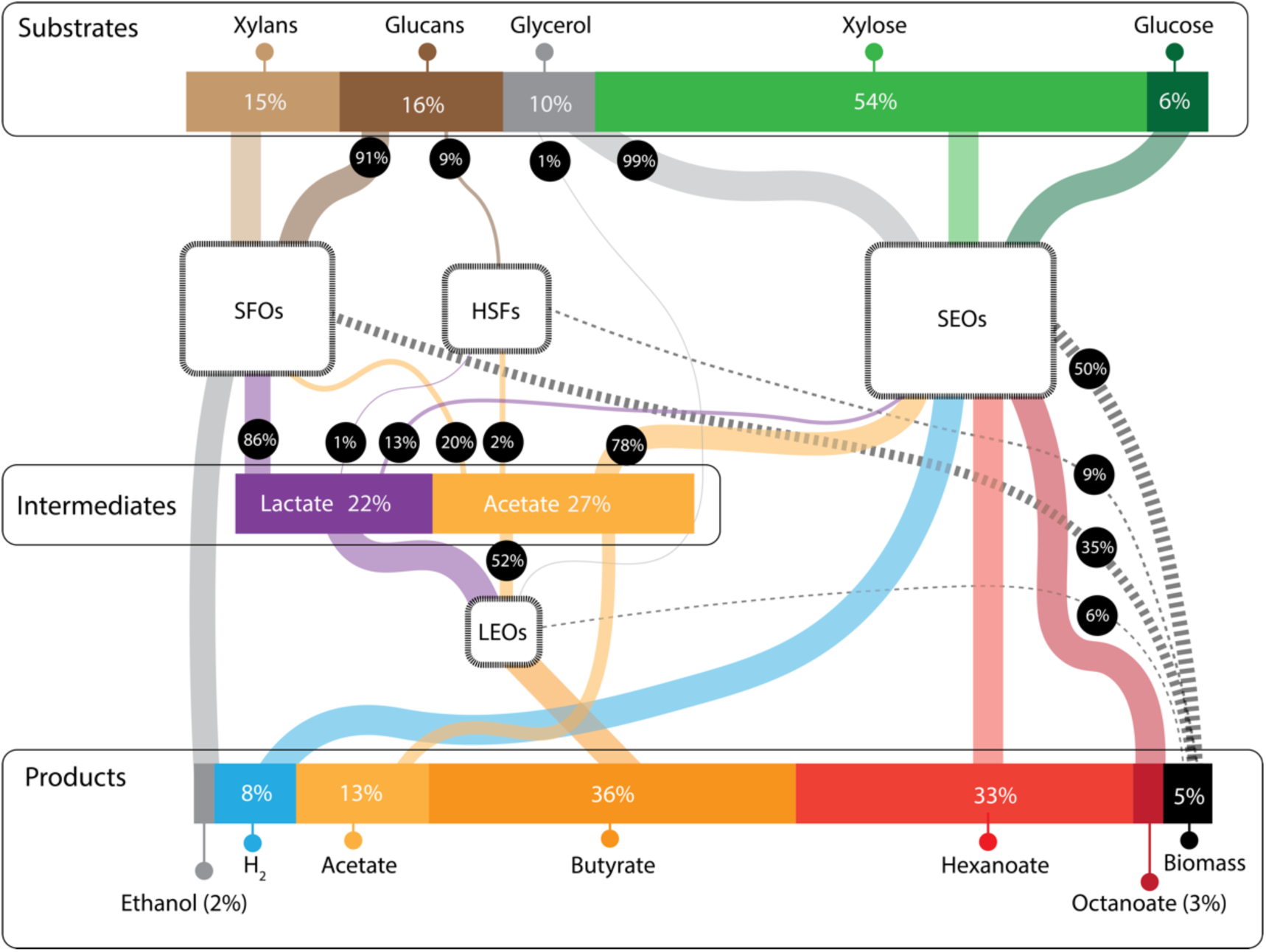
Predicted substrate flow through a mixed culture fermentation producing a mixture of MCFA from a complex feedstock using iFermGuilds789. Values were obtained using pFBA while constraining the products according to observed bioreactor performance. The relative abundance of each functional guild was assumed based on reactor DNA abundance. The relative amount of eeq in the substrates, products, and intermediates are also provided. Line width indicates the relative amount of each substrate consumed and product created by each of the functional guilds. H_2_ was not measured, so for these simulations it was assumed that the difference between eeq in the substrate and eeq in the measured soluble and insoluble products represented H_2_ production. In this simulation, 22% of the eeq in conversion residue was predicted to be directed to lactate as an intermediate and 27% to acetate.

### Predicted routes of sugar and lactate elongation

The above prediction that essentially all MCFAs were produced by SEOs consuming xylose, glycerol, and glucose, while LEOs produced C4 from lactate and C2 were different than the predictions made earlier based on transcriptomic analyses alone.^15^ To gain further insights into SEO and LEO metabolism, we investigated the specific reactions predicted for both guilds by iFermGuilds789. SEOs are predicted to produce pyruvate via glycolysis. Some pyruvate is predicted to be converted to lactate, while most is converted to acetyl-CoA via pyruvate flavodoxin oxidoreductase (PFOR, **Fig 8A**). iFermGuilds789 also predicted that acetyl-CoA is derived from the phosphoketolase pathway (PKETX, **Fig 8A**). Acetyl-CoA is predicted to be directed to reverse β-oxidation, but a portion is also predicted to be hydrolyzed by an acyl-CoA hydrolase. Acyl-CoA hydrolase enzymes are also predicted as terminal enzymes of reverse β-oxidation rather than energy-conserving CoA transferase enzymes (CoAT). SEOs are further predicted to use Hyd1 for hydrogen production, to generate IMF via RNF1, and produce ATP via ATP synthase (ATPS4r) (**Fig. 8A**).

**Figure 8.**
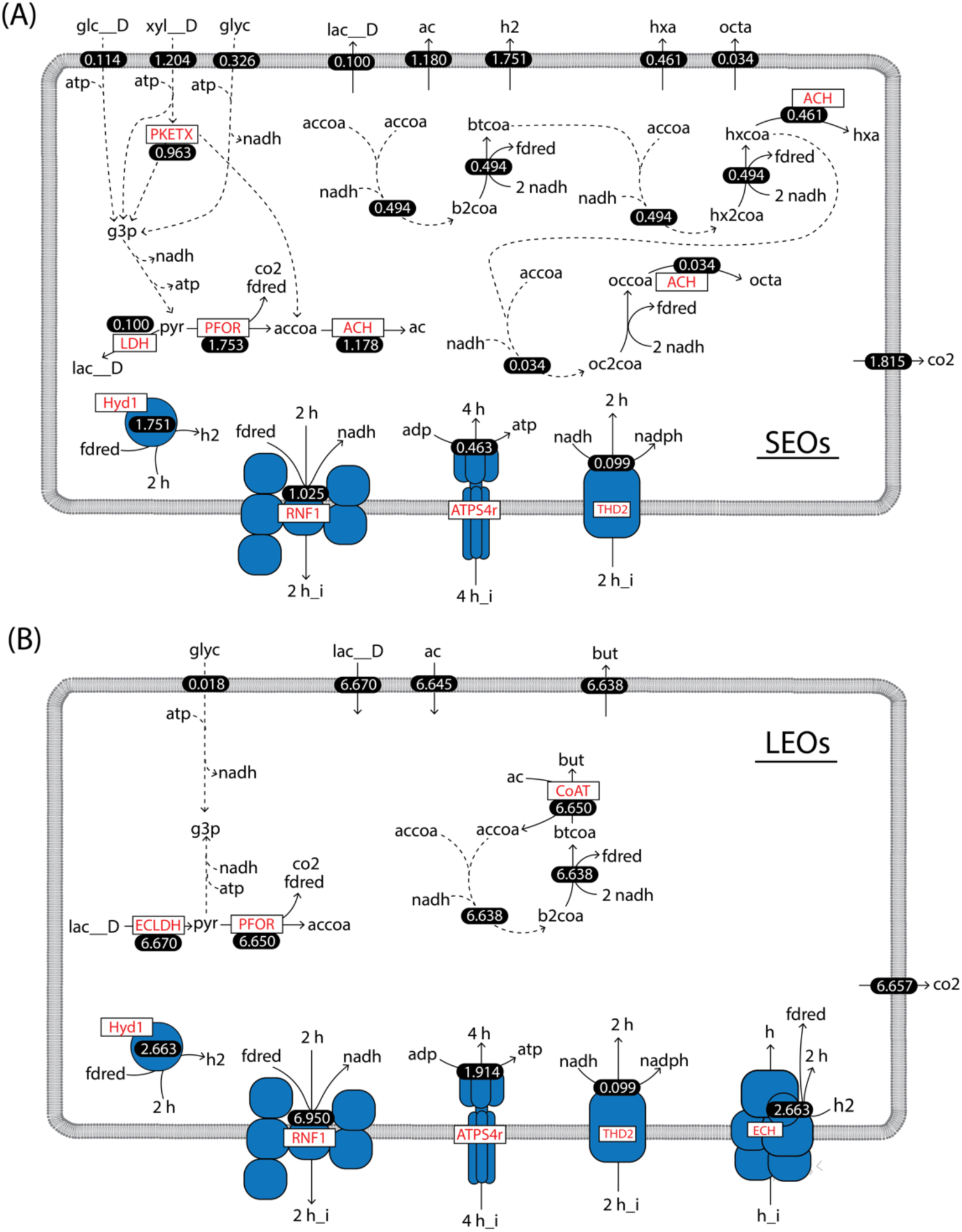
Predicted chain elongation metabolism by SEOs and LEOs using iFermGuilds789. Predicted fluxes values are shown as white numbers and have units of mmol gDCW^-1^ hr^-1^. (A) SEOs are predicted to produce all of the hexanoate (hxa) and octanoate (octa) in the community and use an acyl-CoA hydrolase (ACH) reaction as the terminal step of reverse β-oxidation. (B) LEOs are predicted to produce all of the butyrate in the community while using a CoA transferase (CoAT). LEOs are predicted to recycle all of the H_2_ that they produce via an energy conserving hydrogenase.

LEOs are predicted to convert lactate to pyruvate via an electron-confurcating lactate dehydrogenase (**Fig. 8B**, ECLDH), and pyruvate is predicted to form acetyl-CoA via PFOR. All of the acetyl-CoA is predicted to undergo reverse β-oxidation and C2 is incorporated by a CoAT that transfers the CoA from butyryl-CoA to acetyl-CoA (**Fig. 8B**). LEOs are predicted to consume equimolar concentrations of C2 and lactate, which was previously predicted to favor C4 production (**Fig. 4**). Further, LEOs are not predicted to produce H_2_. iFermGuilds789 predicted that H_2_ produced by Hyd1 is re-consumed via a proton-translocating energy conserving hydrogenase that produces reduced ferredoxin (ECH, **Fig. 8B**). This ferredoxin augments that produced by PFOR and is predicted to be used for lactate consumption by ECLDH and the RNF complex. The use of an energy conserving hydrogenase for ferredoxin production from H_2_ has been demonstrated previously.^40^

While both SEOs and LEOs are predicted by iFermGuilds789 to use reverse β-oxidation to elongate acetyl-CoA, the energy-conserving and redox-balancing reactions needed to support this process are predicted to differ. While SEOs are predicted to use an acyl-CoA hydrolase as the terminal enzyme of reverse β-oxidation, LEOs are predicted to use a CoAT which conserves energy by activating C2 to acetyl-CoA, which would otherwise require ATP. In the previous metatranscriptomic analyses of these community members,^15, 39^ we found that both SEOs and LEOs expressed genes for both acyl-CoA hydrolases and CoAT. The specificity of these predicted chain elongating enzymes for different substrates, however, is not known. The iFermGuilds789 model predictions reinforce the potential importance of terminal enzymes in reverse β-oxidation in determining the final product length; specifically, it predicts that all C4 is produced using CoAT and all MCFAs are using acyl-CoA hydrolase.

Furthermore, SEOs are predicted by iFermGuilds789 to maintain redox balance by producing H_2_ with Hyd1, while LEOs are predicted to recycle all H_2_ internally using both Hyd1 and ECH. Previous metatranscriptomic analyses showed that both LEOs and SEOs expressed genes for both Hyd1 and ECH.^15^ This agrees with iFermGuilds789 predictions for LEOs, but not with SEOs that are predicted to only use Hyd1. When constraining iFermGuilds789 to require SEOs to use ECH, there are no changes in interspecies metabolite transfer (**Supplementary Data File 5)**, but SEOs are predicted to generate additional ATP via ATP synthase while reducing flux through other energy-conserving pathways to achieve the same growth rate (**Supplementary Data File 5**). These findings suggest that SEOs and LEOs have multiple methods for conserving energy and maintaining redox balance and this metabolic flexibility can maintain the same overall transformation of complex substrates to products. This metabolic flexibility may reflect ecological advantages for fermenting organisms, but are also challenges for the accurate predictions of MCFA production, which may strongly depend on the specific organisms present in the microbial community.

## DISCUSSION

Under anaerobic conditions, a diverse set of mixed microbial communities catalyze the transformation of organic substrates to a variety of products. For over a century, anaerobic microbial communities have been harnessed to recover energy, in the form of methane-rich biogas, from complex organic matter in anaerobic digestion.^2^ Some fermentation metabolisms are well understood, and are used by genetically tractable pure cultures that can be analyzed, modeled and improved for use in industrial applications.^41, 42^ In contrast, the activities of mixed microbial communities have been harder to analyze, predict or redirect to improve substrate utilization or product yields. The ability to harness the metabolic activity of mixed microbial communities to produce alternative fermentation products is limited since metabolic and kinetic models that accurately predict the distribution of fermentation products are only starting to emerge.^11-13^ To address this knowledge gap, we constructed metabolic models of mixed culture fermentations with the objective of gaining greater insights into how to increase MCFA production by mixed microbial communities. We focused modeling the production of C6 and C8, linear medium-length MCFAs of industrial value that can be produced from a variety of organic residues.^3^ To build and test these models we took advantage of prior metagenomic, metatranscripotomic, and metabolomic analyses of a mixed community that produces C6 and C8 fatty acids from a lignocellulosic refinery residue.^15, 39^

The predictions of our models generated several hypotheses that should be explored with further analyses. First, the single unit iFermCell215 predicted production of MCFAs from ethanol as a sole source of electrons and carbon (**Fig. 4A-D**), and this is supported by recent experiments with mixed cultures.^33^ More work is needed to determine bioreactor conditions that promote elongation of ethanol as a sole substrate or with low C2:EtOH ratios. The iFermCell215 simulations predict a threshold of 0.56 mol C2 per mol ethanol below which C8 production benefits from C2 co-utilization, and above which C8 production starts to decrease and to be replaced by production of shorter acids. This shift in product profile was predicted by iFermCell215 to be associated with reversal in the direction of enzymatic activities related to energy conserving metabolism, an observation that still lacks experimental verification.

Second, even though lactate is a known substrate for chain elongation,^9^ iFermGuilds789 simulation results suggest that, in this mixed community, lactate may be converted to C4 instead of C6 or C8 **(Fig. 7)**. Although at lower C2 to lactate ratios iFermCells215 simulations predicted that this ratio was not as influential on determining the product distribution as the C2:EtOH ratio, it also predicted that excess C2 could result in C4 production instead of C6 or C8 from both intermediate metabolites (**Fig. 4**). The iFermGuilds789 simulation predicted co-utilization of C2 and lactate at a molar ratio of 1.0 (**Fig. 8B**), which according to the iFermCell215 simulations (**Fig. 4**) this is already an excess C2 condition predicted to yield C4 as the main fermentation product. These results have the important implication that efforts directed to construct synthetic communities or control self-assembled communities for production of MCFA should focus on minimizing C2 production and accumulation.

Third, iFermCell215 modeling results suggest that controlling product formation from lactate may not be possible by controlling C2-to-lactate ratios (**Fig. 4E-G**), whereas C2:EtOH ratios suggests very specific targets to maximize C8 production (**Fig. 4A-D**). This suggests that for MCFA production from complex organic substrates, in addition to targeting the reduction of C2 production, efforts should also be directed to creating conditions in which ethanol instead of lactate is the key intermediate metabolite for MCFA production. However, this may not be achievable in self-assembled communities, and since simulations predict that the products of lactate fermentation are dependent on specific metabolic features of LEOs (**Fig S8**), it is important to continue the discovery of the natural diversity of lactate fermentation metabolisms, which eventually could help identify preferred LEOs for the development of novel strategies to promote MCFA production when lactate is the key intermediate metabolite.

Fourth, iFermGuilds789 predicts that enzyme affinity plays a major role in determining the final products of reverse β-oxidation and mixed culture fermentation (**Fig 6D**). Additional work is needed to assess the affinity of reverse β-oxidation, acyl-CoA hydrolase, and CoA transferase enzymes for different chain lengths to identify ideal microorganisms or engineer microorganisms to improve MCFA production. Past studies have identified and engineered acyl-ACP thioesterase enzymes (the terminal enzyme of fatty acid synthesis via the fatty acid biosynthesis pathway) with high affinity for C8.^43-45^ Similar studies should be performed with acyl-CoA hydrolase and acyl-CoA transferase enzymes to characterize and improve the terminal enzymes of reverse β-oxidation.

Since many microbes in natural and self-assembled fermentation communities remain uncultured, our knowledge and ability to predict important metabolic networks relies largely on information from metagenomics, metatranscriptomics, and metabolomics analyses to build the functional activities in each guild.^15, 46^ We show in this work that metabolic modeling offers additional insight that is not gained by these analyses, but that is critical in the evaluation of the microbial communities and in developing hypothesis of how to direct metabolism towards a specific set of fermentation products. From prior analyses,^15^ we anticipated that both SEOs and LEOs participated in MCFA production, but the iFermGuilds789 simulation of interspecies metabolite flux (**Fig. 7**) predicted that to achieve the observed growth rate of the microbial community, C6 and C8 were produced solely by SEOs, while LEOs were responsible for production of C4. This difference between predictions based on metatranscriptomic and simulation analyses can be tracked to the importance of co-utilization of fermentation products (i.e., C2 and lactate), which is easier to explore using metabolic models.

iFermCell215 and iFermGuilds789 are examples of first-generation predictive models of a mixed culture fermentation. Their utility can be improved by obtaining additional knowledge of metabolic pathways involved in fermentation through studying individual microbes within each functional guild. Further effort is needed to simulate poorly appreciated conditions that shape community structure, such as the toxicity of fermentation products on the specific guilds or the contribution(s) of low abundance microbes to the overall activity of the mixed culture fermentation. Additionally, computational methods to predict the abundance of specific guilds, rather than relying on relative abundances as a model input, are needed. Efforts to assemble such models have begun for the human gut microbiome,^47, 48^ and as these approaches are applied to mixed microbial fermentations, their benefits to the production of MCFA and other chemicals of societal importance will continue to improve.

## ACKNOWLEDGMENTS

This work was funded by the DOE Great Lakes Bioenergy Research Center (DOE BER Office of Science DE-SC0018409). M.J.S. was supported by the National Science Foundation Graduate Research Fellowship Program under grant No. DGE-1256259 and by the University of Vermont College of Engineering and Mathematical Sciences. J.J.H. was supported by the National Institute of Food and Agriculture, U.S. Department of Agriculture, under award number 2016-67012-24709. The authors extend our gratitude to Robert Aidan Campbell for assistance with Python coding.

