## Supplementary Information for "Diagnosing and predicting mixed culture fermentations with unicellular and guild-based metabolic models"

**Supplementary Text 1: Examples of organisms for each functional guild**

SEOs include organisms that utilize sugars and elongate intermediate products through reverse β-oxidation, such as *Megasphaera* and *Caproicproducens*.[1, 2] SFOs include genera such as *Lactobacillus* and *Bifidobacterium*, which can ferment hexoses and pentoses to lactate, acetate and ethanol.[3, 4] HSFs include organisms that generate $H_2$ while producing fermentation products, such as members of the *Coriobacteriaceae* family.[5] LEOs include organisms that perform reverse β-oxidation with lactate, such as *Pseudoramibacter*[6] and *Ruminococcaceae* bacterium *CPB6.*[7] EEOs, such as *Clostridium kluyveri,*[8] perform reverse β-oxidation with ethanol. HAOs produce acetate from $H_2$ and $CO_2$ and include species within the *Clostridium*, *Acetobacterium*, *Eubacterium,* and *Blautia.*[9-11]

**Supplementary Text 2: Formation of a simplified biomass equation**

Simulating biomass growth requires a stochiometric equation accounting for required biomass precursors. Bacteria differ in their cellular composition, but gram positive bacteria are often abundant in fermentation bioreactors [7, 12-15]. We therefore assumed a biomass composition similar to the well-studied gram positive bacterium *B. subtilis*, which contains proteins, nucleic acids, lipids, peptidoglycan, and lipoteichoic acid [16, 17]. Stoichiometric coefficients for each precursor were calculated based on established pathways for synthesis of biomass components [18]. We converted the stoichiometry for biomass production into 17 precursors contained in the iFerment215 and iFermGuilds789: α-ketoglutarate (aKG), oxaloacetate (OAA), glucose-6-phosphate (G-6-P), glucose-3-phosphate (G-3-P), 3-phosphoglycerate (3-PG), pyruvate, phosphoenolpyruvate, acetyl-CoA, erythrose-4-phosphate, ribulose-5-phosphate, ATP, NADPH, NAD+, ammonium, sulfate, and $H_2O$.

**Protein biosynthesis**

Based on established amino acid biosynthesis pathways (**Figs. S1 and S2**), we calculated the stoichiometry for synthesis of each amino acid. The final stoichiometry for each amino acid synthesis pathway is provided in **Supplementary Data File 1**.

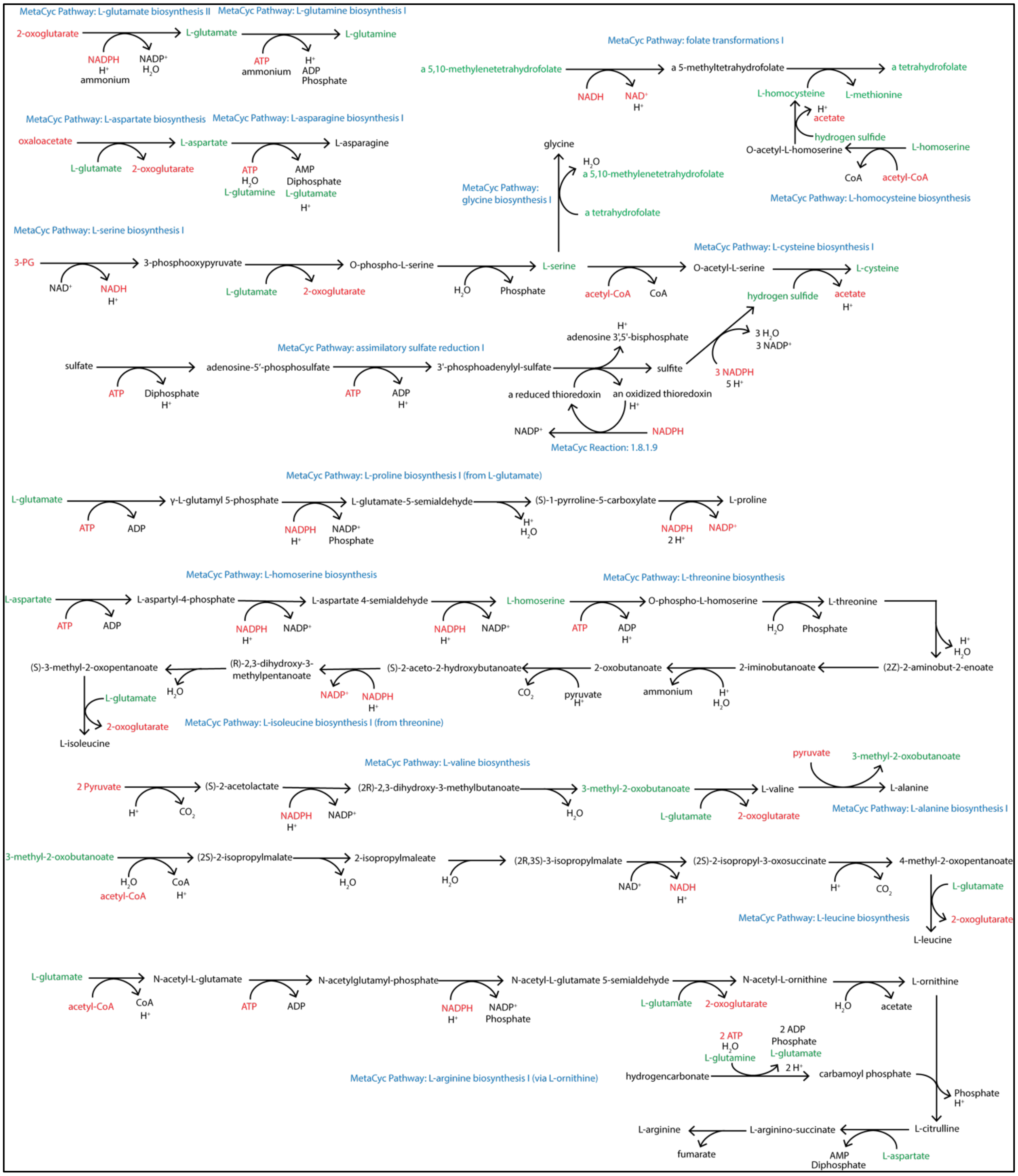


**Fig S1.** Metabolic pathways for synthesis of L-glutamate, L-glutamine, L-aspartate, L-asparagine, L-serine, glycine, L-cysteine, L-proline, L-threonine, L-isoleucine, L-valine, L-alanine, L-Leucine, and L-arginine.

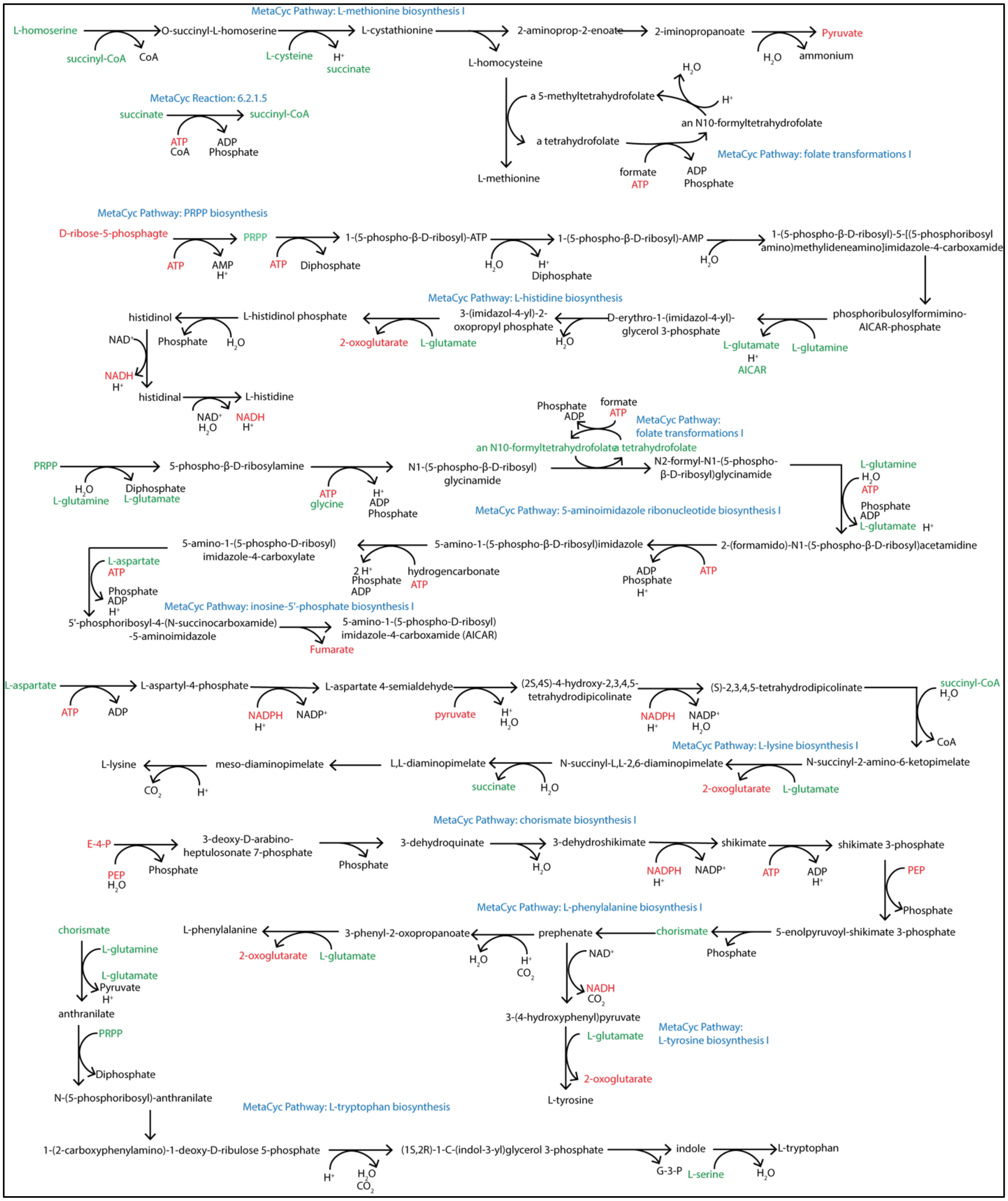


**Fig S2.** Metabolic pathways for synthesis of L-methionine, L-histidine, L-Lysine, L-phenylalanine, L-tyrosine, and L-tryptophan.

### Additional ATP consumption for protein synthesis

The synthesis of proteins from amino acids requires the equivalent of 4 mol ATP per mol amino acid incorporated into a protein. Aminoacyl-tRNA synthetase uses the equivalent of 2 mol ATP to load an amino acid into a tRNA,[19] and 2 GTP are hydrolyzed by elongation factors (creating 2 GDP) for each peptide bond formed.[20] Therefore, in addition to the ATP required to synthesize the amino acid, it was assumed that 4 mol ATP are required for each mol of amino acid incorporated into biomass.

### Nucleic acid synthesis

RNA and DNA are formed by the polymerization of four different nucleoside triphosphates (NTPs: ATP, GTP, UTP, CTP) and four different deoxy-nucleoside triphosphates (dNTPs: dATP, dGTP, dTTP, dCTP), respectively. The synthesis pathways assumed for these nucleotides are shown in **Fig S3**.

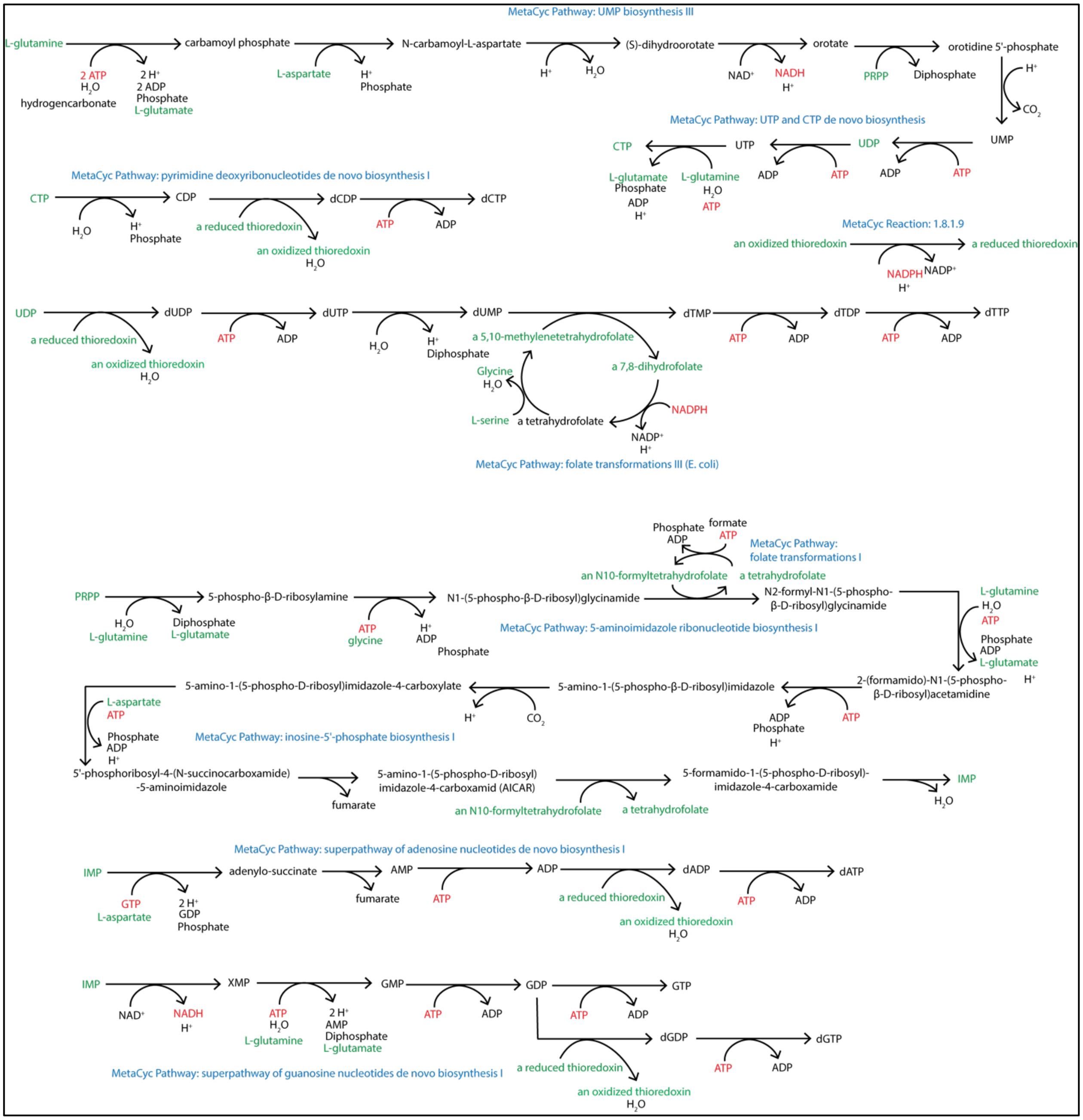


**Fig. S3. Metabolic pathways for nucleotide synthesis**

### Lipid Biosynthesis

We assumed phosphotidylethanolamine as a representative lipid, which is the most abundant lipid reported in *B. subtilis* [21]. Palmitate (C16:0), which is representative of the average size of fatty acid contained in *B. subtilis* lipids [21], was assumed to be the sole fatty acid incorporated into phosphotidylethanolamine (**Fig. S4**).

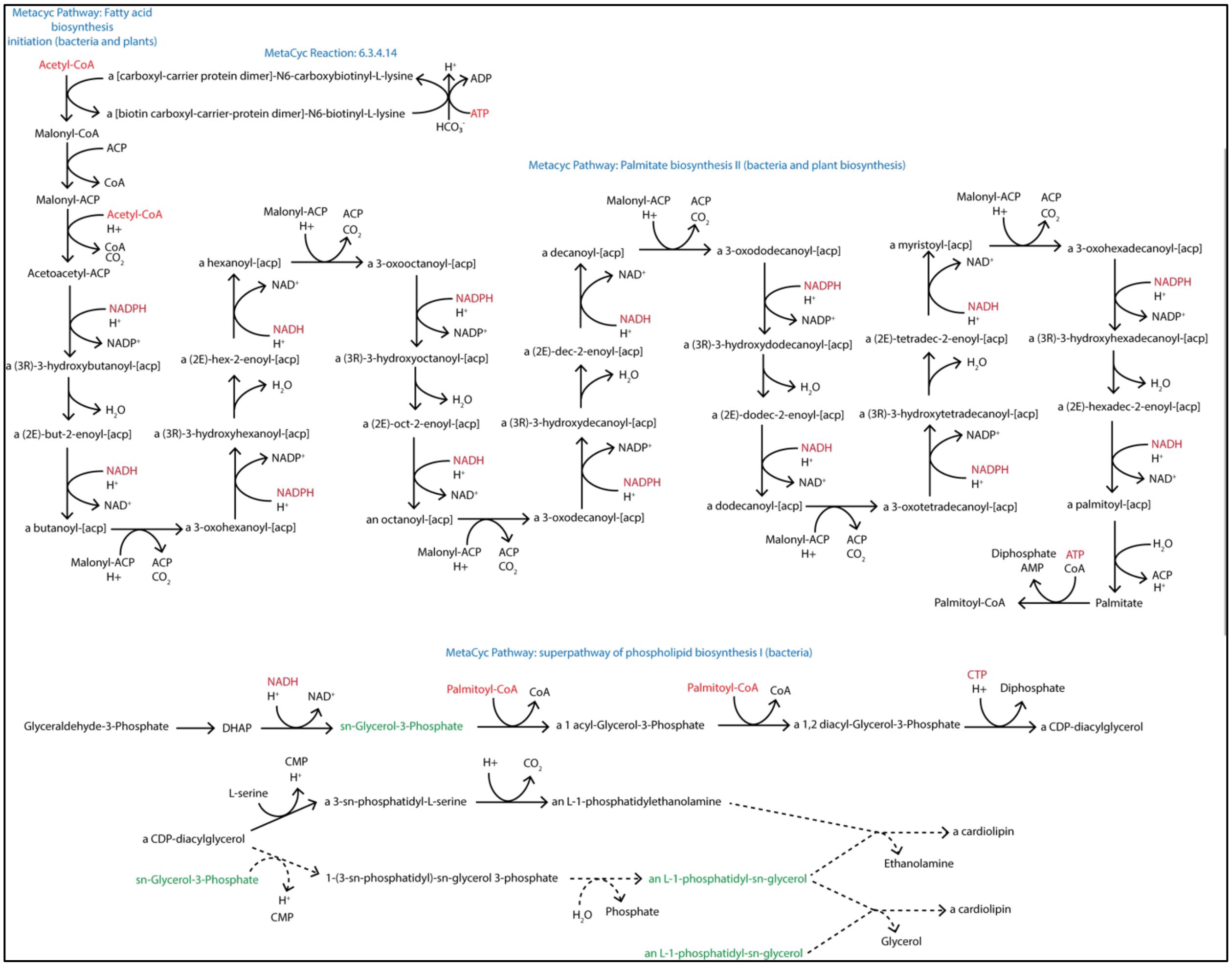


**Fig. S4.** Metabolic pathways for phospholipid biosynthesis

### Peptidoglycan biosynthesis

We assumed peptidoglycan is constructed of subunits containing N-acetylglucosamine, N-acetylmuramic acid, and a pentapeptide consisting of 1 L-alanine, D-glutamine, L-Lysine, and two

D-alanine. We also assumed that the peptidoglycan subunits contain di-trans,octa-cis-undecaprenyl phosphate. Key metabolic intermediates of peptidoglycan synthesis include UDP-N-acetyl-α–glucosamine, isopentenyl diphosphate, and di-trans,octa-cis-undecaprenyl phosphate (**Fig. 5**).

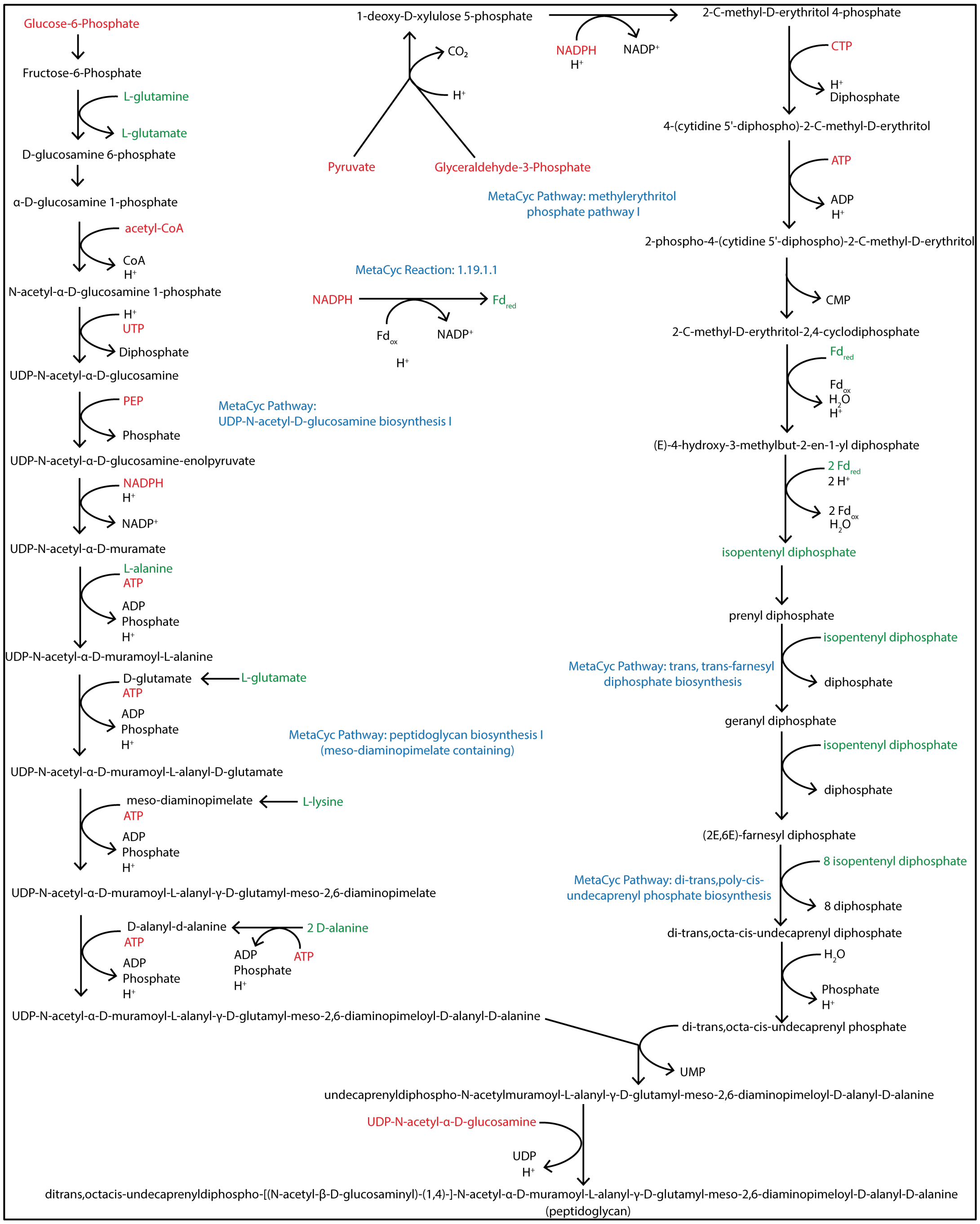

**Fig. S5.** Metabolic pathways for peptidoglycan synthesis

**Lipoteichoic acid biosynthesis**

Lipoteichoic acid (LTA) is a major component of gram positive cell walls consisting of a glycerol backbone and varying sugar units that is attached to the cell membrane via glycolipids. Type 1 LTA is widespread and found in many Firmicutes [22, 23]. Production of LTA relies on several precursors (**Fig. 6**), including G-6-P, an L-1-phosphaidyl-sn-glycerol (an intermediate of phospholipid synthesis, **Fig. 5**), UDP-N-acetyl-α–glucosamine , and D-alanine.

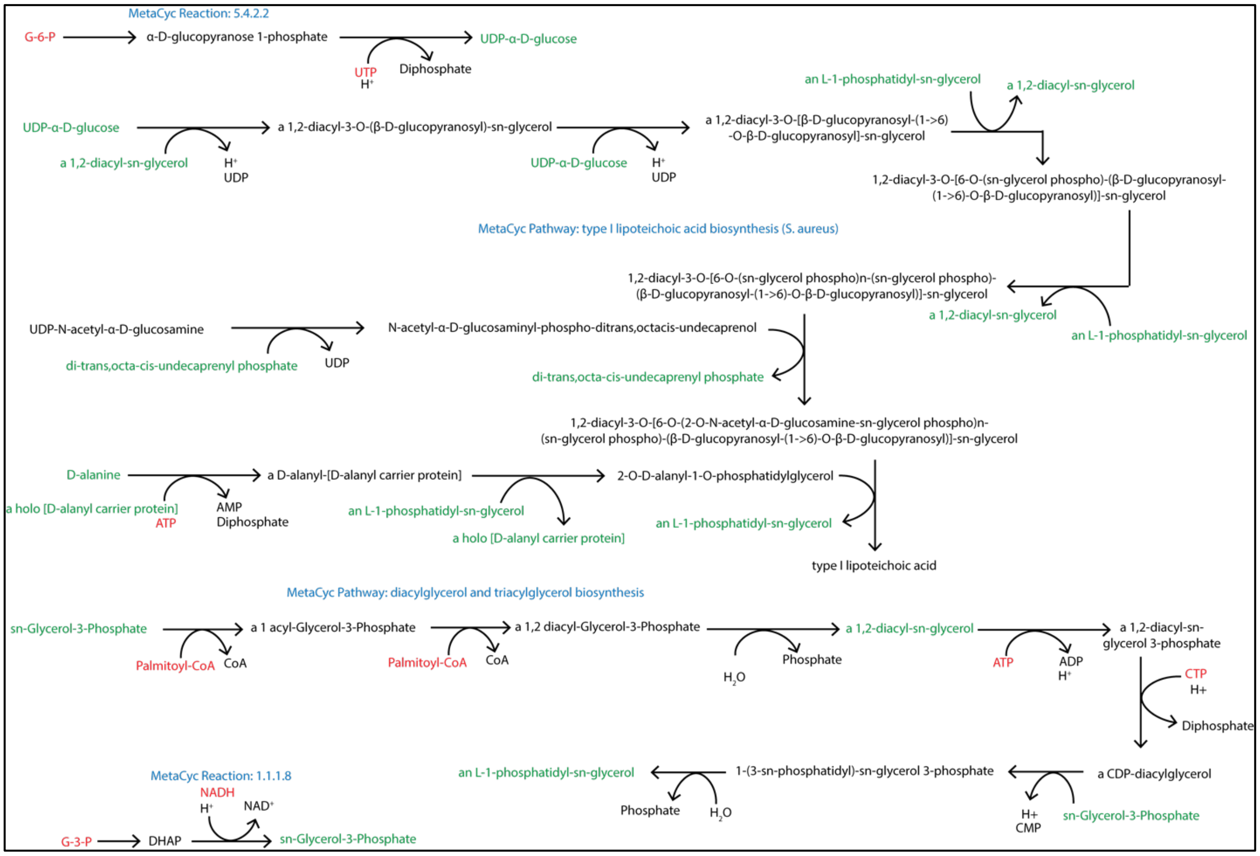


**Fig. S6.** Metabolic pathways for lipoteichoic acid synthesis

**Formation of the biomass equation**

After creating stoichiometric matrices for each biomass component (**Supplementary Data File 1**), we created a biomass equation assuming a cellular composition based on the well-studied gram positive bacterium *Bacillus subtilis*. We assumed biomass composition as a percent of cell dry weight based on previous biomass analyses.[17] For the biomass equation, we considered protein, RNA, DNA, lipids, peptidoglycan, and teichoic acid.

**Table S1. Assumed biomass composition**

| **Biomass component** | **Percent of dry cell weight (%)** | **Molecular weight (g $mol^{-1}$)** | **mmol $gDCW^{-1}$** |
|---|---|---|---|
| **Protein** | **55.6** | | |
| L-glutamate | 4.0 | 146.1 | 0.27 |
| L-glutamine | 4.0 | 146.1 | 0.27 |
| L-aspartate | 2.1 | 132.1 | 0.16 |
| L-asparagine | 2.1 | 132.1 | 0.16 |
| L-serine | 2.4 | 105.1 | 0.23 |
| L-cysteine | 0.7 | 121.2 | 0.06 |
| L-proline | 1.9 | 115.1 | 0.17 |
| L-threonine | 2.3 | 119.1 | 0.19 |
| L-isoleucine | 3.7 | 131.2 | 0.28 |
| L-valine | 3.8 | 117.1 | 0.32 |
| L-alanine | 2.5 | 89.1 | 0.28 |
| L-leucine | 4.8 | 131.2 | 0.37 |
| L-arginine | 3.6 | 175.2 | 0.21 |
| L-methionine | 1.8 | 149.2 | 0.12 |
| Glycine | 3.2 | 75.1 | 0.43 |
| L-histidine | 1.3 | 155.2 | 0.08 |
| L-lysine | 5.0 | 147.2 | 0.34 |
| L-phenylalanine | 3.1 | 165.2 | 0.19 |
| L-tyrosine | 2.1 | 181.2 | 0.12 |
| L-tryptophan | 1.2 | 204.2 | 0.059 |
| **RNA** | **6.9** | | |
| AMP (ATP) | 1.8 | 345.2 | 0.052 |
| GMP (GTP) | 2.4 | 361.2 | 0.066 |
| CMP (CTP) | 1.3 | 321.2 | 0.040 |
| UMP (UTP) | 1.4 | 322.2 | 0.043 |
| **DNA** | **2.7** | | |
| dAMP (dATP) | 0.79 | 329.2 | 0.024 |
| dGMP (dGTP) | 0.62 | 345.2 | 0.018 |
| dCMP (dCTP) | 0.55 | 305.2 | 0.018 |
| dTMP (dTTP) | 0.76 | 320.2 | 0.024 |
| **Lipid** | **8.0** | | |
| Phosphotidylethanolamine (C 16:0 sidechains) | 8.0 | 480.1 | 0.17 |
| **Peptidoglycan** | **23.6** | | |
| Peptidoglycan (meso-diaminopimelate containing) | 23.6 | 1,916.2 | 0.12 |
| **Lipoteichoic acid** | **3.2** | | |
| Type 1 Lipoteichoic acid | 3.2 | 2,291.4 | 0.014 |

We then determined the mmol of each precursor needed to produce one gram of dry cell weight (gDCW) by multiplying the stoichiometryby the mmol gDCW$^{-1}$ (**Supplementary Data File 1**). This yielded the stoichiometry shown in the **Supplementary Data File 1 – Stoichiometric Matrix**. Minor modifications were made to the precision of calculated stoichiometric coefficients (**Supplementary Data File 1 – Biomass Equation**) to allow the model to achieve mass balance with biomass coproducts. The resulting biomass equation, based on 1 mmol of biomass having a mass of 1 gDCW, is shown below.

1.17 aKG + 2.06 OAA + 0.26 G-6-P + 1.58 G-3-P + 1.31 3-PG + 4.33 Pyruvate + 0.92 PEP + 3.06 Acetyl-CoA + 0.40 E-4-P + 0.35 R-5-P + 36.0 ATP + 19.39 NADPH + 1.10 $NAD^+$ + 8.62 Ammonium + 0.21 Sulfate + 7.57 $H_2O$ $\leftrightarrow$ 1 Biomass + 0.37 Fumarate + 0.43 Acetate + 0.29 Formate + 1.6 $NAD^+$ + 19.4 $NADP^+$ + 34.6 ADP + 1.4 AMP + 31.88 Pi + 4.74 PPi + 1.10 NADH + 3.54 $CO_2$ + 10.13 $H^+$ + 3.06 CoA

This stoichiometry was added for each biomass reaction in iFerment215 and iFermGuilds789. An example from iFermCell215 is shown below.

```
reaction.add_metabolites({akg_CELLc:        -1.17,
                          oaa_CELLc:        -2.06,
                          g6p_CELLc:        -0.26,
                          g3p_CELLc:        -1.58,
                          _3pg_CELLc:       -1.31,
                          pyr_CELLc:        -4.33,
                          pep_CELLc:        -0.92,
                          accoa_CELLc:      -3.06,
                          e4p_CELLc:        -0.40,
                          r5p_CELLc:        -0.35,
                          fum_CELLc:         0.37,
                          ac_CELLc:          0.43,
                          for_CELLc:         0.29,
                          atp_CELLc:        -36.0,
                          nadph_CELLc:     -19.39,
                          nadh_CELLc:        1.10,
                          nh4_CELLc:        -8.62,
                          h_CELLc:          10.13,
                          adp_CELLc:         34.6,
                          pi_CELLc:         31.88,
                          ppi_CELLc:         4.74,
                          amp_CELLc:          1.4,
                          co2_CELLc:         3.54,
                          h2o_CELLc:        -7.57,
                          coa_CELLc:         3.06,
                          nad_CELLc:        -1.10,
                          nadp_CELLc:       19.39,
                          so4_CELLc:        -0.21,
                          BIOMASS_CELLe:        1,
                          })
```

**Supplementary Text 3: Transport Constraints used in metabolic models**

To describe bioenergetic requirements for transport, we considered contributions of the electrochemical, pH, and concentration gradients across the cell envelope (**Eq. S1**).[24] This predicts the energy required to transport a product from the interior of the cell to the extracellular space as represented by Equation 1, where R is the ideal gas constant (8.315 x $10^{-3}$ kJ $mol^{-1}$ $K^{-1}$), T is the temperature (K), $[X]_{ext}$ is the concentration of product in the bulk liquid (M), $[X]_{int}$ is the concentration of product inside a cell (M), $\Delta\psi$ is the difference in electric potential across the membrane (mV), $c_j$ is the net charge transported from inside to outside the cell, F is Faraday's constant (0.096485 kJ $mV^{-1}$ $mol^{-1}$), $h_j$ is the number of protons transported across the membrane, and $\Delta pH$ (standard pH units) is the difference in pH between the intracellular and extracellular environments. We assumed an intracellular pH of 7.0 and an extracellular pH of 5.5 for all modeling scenarios, consistent with experimentational observations of mixed culture fermentation reactors[25] and previously published models.[26] Based on these conditions, we modeled the transport of unprotonated forms of products based on the $pK_A$ values of carboxylic acid end products. Based on previous metabolic modeling approaches,[27] we estimated the $\Delta\psi$ as shown in **Equation S2**.

$$\Delta G_t = RTln\left(\frac{[X]_{ext}}{[X]_{int}}\right) + \Delta\psi c_j F - 2.3 h_j RT \Delta pH \qquad \textbf{(Eq. S1)}$$

$$\Delta\psi = 33.3\Delta pH - 143.33 \qquad \textbf{(Eq. S2)}$$

As such, the $\Delta\psi$ was equal to -93.4mV which is similar to potentials measured in other fermenting cultures.[28] We further constrained intracellular product concentrations to 10 mM, consistent with previous studies[26] and extracellular product concentrations to experimentally measured values obtained from the bulk liquid of a MCFA-producing mixed culture fermentation reactor.[29] These bioenergetic calculations were then used to impose additional constraints on individual models, requiring that ATP hydrolysis for transport of end products be deducted from the gross ATP produced by the cell. This results in a net ATP yield that is a proxy for cell growth.

**Table S2.** Summary of metabolites that are consumed or produced in the metabolic models used in this study. Chemical formulas are provided for the predominant form at a pH of 7.0.

| **Name** | **Abbreviation** | **Formula** | **eeq mol$^{-1}$** | **$\Delta G_f^0$ (kJ mol$^{-1}$)** |
|---|---|---|---|---|
| Glucose | glc | $C_6H_{12}O_6$ | 24 | -913 |
| Xylose | xyl | $C_5H_{10}O_5$ | 20 | -753 |
| Glucan (stachyose) | glc4 | $C_{24}H_{48}O_{24}$ | 96 | -2,906 |
| Xylan (tetra-arabinofuranoside) | xyl4 | $C_{20}H_{34}O_{17}$ | 90 | -2,265 |
| Glycerol | glyc | $C_3H_8O_3$ | 14 | -486 |
| Ethanol | etoh | $C_2H_6O$ | 12 | -182 |
| Lactate | lac | $C_3H_6O_3^-$ | 12 | -515 |
| Formate | for | $C_1H_1O_2^-$ | 2 | -351 |
| Acetate | ac | $C_2H_3O_2^-$ | 8 | -369 |
| Propionate | ppa | $C_3H_5O_2^-$ | 14 | -356 |
| Butyrate | but | $C_4H_7O_2^-$ | 20 | -353 |
| Valerate | pta | $C_5H_9O_2^-$ | 26 | -343 |
| Hexanoate | hxa | $C_6H_{11}O_2^-$ | 32 | -336 |
| Heptanoate | hpta | $C7H_{13}O_2^-$ | 38 | -329 |
| Octanoate | octa | $C8H_{15}O_2^-$ | 44 | -322 |
| Hydrogen | h2 | $H_2$ | 2 | 0 |
| Hydrogen ions | h | $H^+$ | 0 | 0 |
| Carbon dioxide | co2 | $CO_2$ | 0 | -386 |
| Water | h2o | $H_2O$ | 0 | -237 |

**Supplementary Text 4: Ability of iFermCell215 to predict known fermentation pathways**

To test the predictive ability of iFermCell215, we modeled several glucose fermentation pathways. By setting the glucose uptake flux and constraining iFermCell215 to produce defined products (**Table S2**), we assessed if the model could recapitulate known product formation stoichiometries. We also modeled conditions that produce C6 or C8 as sole products from glucose, which to our knowledge, has not been experimentally observed. Under each condition, we evaluated potential ATP yields and the efficiency of carbon transformation (**Table S3**).

iFermCell215 simulated known glucose fermentation stoichiometries, including homofermentations to lactate,[30] ethanol,[31] and butyrate,[32] and heterofermentative production of acetate and propionate,[33] acetate and lactate,[34] and acetate and $H_2$.[35] This single unit model predicted that conversion of glucose to butyrate could result in a higher ATP yield than producing acetate, lactate, or ethanol. This prediction is consistent with existing literature suggesting that butyrate production via reverse β-oxidation can conserve additional energy as ATP.[36, 37] iFermCell215 also predicted that homofermentative production of C6 or C8 (**Table S3**) could increase the ATP yield compared to butyrate production, as proposed previously.[29] These results predict that fermentative pathways can have large impacts on ATP yield. For instance, if the single "organism" modeled by iFermCell215 is only producing lactate, the expected ATP yield is 2.00 mol ATP $mol^{-1}$ glucose, but if C8 could be produced as a single product, the ATP yield would increase to 3.82 mol ATP $mol^{-1}$ glucose, a 91% gain in the yield of this cellular energy source. The agreement of the predictions of iFermCell215 with known bioreactor stoichiometries[1, 6, 30-35] illustrates the applicability of this model. iFermCell215 offers the simplicity of having all the reactions in a single unit, and therefore, can be easily adapted to simulate the activities of individual microbes by constraining the model to using only subsets of the known pathways.

**Table S3.** Glucose fermentation pathways in iFermCell215 and their predicted ATP yields

| Description | Example Organism | iFermCell215 Predicted Reaction | ATP Yield | $\Delta G^{0'}$ mol$^{-1}$ ATP | Carbon Capture |
|---|---|---|---|---|---|
| Homofermentation to Lactate | *Lactobacillus lactis* [30] | $1\ C_6H_{12}O_6 \rightarrow 2\ C_3H_5O_3^- + 2\ H^+$ | 2.00 | -98.6 | 100% |
| Heterofermentation to Lactate and Acetate | *Bifidobacterium animalis* [34] | $1\ C_6H_{12}O_6 \rightarrow 1\ C_3H_5O_3^- + 1.5\ C_2H_3O_2^- + 2.5\ H^+$ | 2.50 | -74.1 | 100% |
| Fermentation to Ethanol | *Escherichia coli K12*[1] [31] | $1\ C_6H_{12}O_6 \rightarrow 2\ C_2H_6O_3^- + 2\ CO_2$ | 2.00 | -74.1 | 66.7% |
| Fermentation to Acetate and Hydrogen | *Thermotoga Maritima* [35] | $1\ C_6H_{12}O_6 \rightarrow 2\ C_2H_3O_2^- + 2\ CO_2 + 2\ H_2 + 2\ H^+$ | 3.00 | -67.7 | 66.7% |
| Fermentation to Propionate and Acetate | *Bacteroides fragilis* [33] | $1\ C_6H_{12}O_6 \rightarrow 1.83\ C_2H_3O_2^- + 0.67\ C_3H_5O_2^- + 0.33\ CO_2 + 2.5\ H^+ + 0.333\ H_2O$ | 3.00 | -103 | 95.5% |
| Butyrate Production | *Corynebacterium glutamicum* [32] | $1\ C_6H_{12}O_6 \rightarrow 1.2\ C_4H_7O_2^- + 1.2\ CO_2 + 1.2\ H^+ + 1.2\ H_2O$ | 3.60 | -84.9 | 80.0% |
| Hexanoate Production | *NA*[2] | $1\ C_6H_{12}O_6 \rightarrow 0.75\ C_6H_{11}O_2^- + 1.5\ CO_2 + 0.75\ H^+ + 1.5\ H_2O$ | 3.75 | -80.9 | 75.0% |
| Octanoate Production | *NA*[2] | $1\ C_6H_{12}O_6 \rightarrow 0.545\ C_8H_{15}O_2^- + 1.636\ CO_2 + 0.545\ H^+ + 1.636\ H_2O$ | 3.82 | -79.6 | 72.6% |

[1] A genetically modified organism was used for the designated process.
[2] Not Applicable. There are no known demonstrations of homo-hexanoate or homo-octanoate fermentation from glucose. Instead, hexanoate and octanoate are often produced with other end products, such as butyrate and acetate.

**Table S4.** Reaction knockouts to meet thermodynamic constraints for scenarios with single substrates. Simulations were used to generate **Fig. 3.**

| **Substrate** | **Flux Sampling Seed** | **Additional Reaction Constraints** | **CobraPy Code** |
|---|---|---|---|
| Glucose | 8211 | Hyd1 irreversible | `model.reactions.HYD1.lower_bound = 0` |
| Xylose | 4844 | Hyd1 irreversible | `model.reactions.HYD1.lower_bound = 0` |
| Glycerol | 3770 | Hyd1 irreversible | `model.reactions.HYD1.lower_bound = 0` |
| Lactate | 561 | Hyd 1 irreversible; FDH irreversible | `model.reactions.HYD1.lower_bound = 0`<br>`model.reactions.FDH.lower_bound = 0` |
| Ethanol | 8108 | Hyd 1 irreversible; FDH irreversible; AckR irreversible | `model.reactions.HYD1.lower_bound = 0`<br>`model.reactions.FDH.lower_bound = 0`<br>`model.reactions.ACKr.lower_bound = 0` |

**Table S5.** Reaction knockouts to meet thermodynamic constraints for scenarios with multiple substrates. Simulations were used to generate **Fig. 3.**

| **Electron Donor** | **Electron Acceptor** | **Flux Sampling Seed** | **Additional Reaction Constraints** | **CobraPy Code** |
|---|---|---|---|---|
| $H_2$ | $CO_2$ | 9814 | Hyd 1 irreversible; Formate export knocked out | `model.reactions.HYD1.lower_bound = 0`<br>`model.reactions.EX_for_CELLe.knock_out()` |
| Ethanol | Acetate | 5350 | Hyd 1 irreversible; FDH irreversible; Acetate kinase irreversible | `model.reactions.HYD1.lower_bound = 0`<br>`model.reactions.FDHlower_bound = 0`<br>`model.reactions.ACKr.lower_bound = 0` |
| Lactate | Acetate | 4100 | Hyd 1 irreversible; FDH irreversible | `model.reactions.HYD1.lower_bound = 0`<br>`model.reactions.FDH.lower_bound = 0` |

### Supplementary Text 5. Impacts of reaction knockouts on predicted products from lactate

To assess the impacts of individual reaction on the predicted products of lactate consumption, we additively knocked out reactions in the iFermCell215 model. After knocking out homoaectogenesis, iFermCell215 predicted a mixture of C2, C4, C6, C8 and $H_2$ as final products (**Fig. S7A**). Without acetate kinase, production of heptanoate (C7) and C8 is predicted (**Fig. S7B**).

Without energy conserving hydrogenases (Ech and HydABC), the model predicted C7 as the most abundant product (**Fig S7B**). Knocking out CoAT results in predicted production of C8, formate, and $H_2$ (**Fig. S7D**) and similar predictions were made after knocking out an NADH-dependent lactate dehydrogenase which required the model to use an electron-confurcating lactate dehydrogenase (**Fig S7E**). Finally, by knocking out Hyd1 (and all H2 production), the model predicted C8 and C1 as the products (**Fig S7F**). In total, these results suggest that excluding homoacetogensis, acetate kinase, and energy-conserving hydorgenases may improve production of C8.

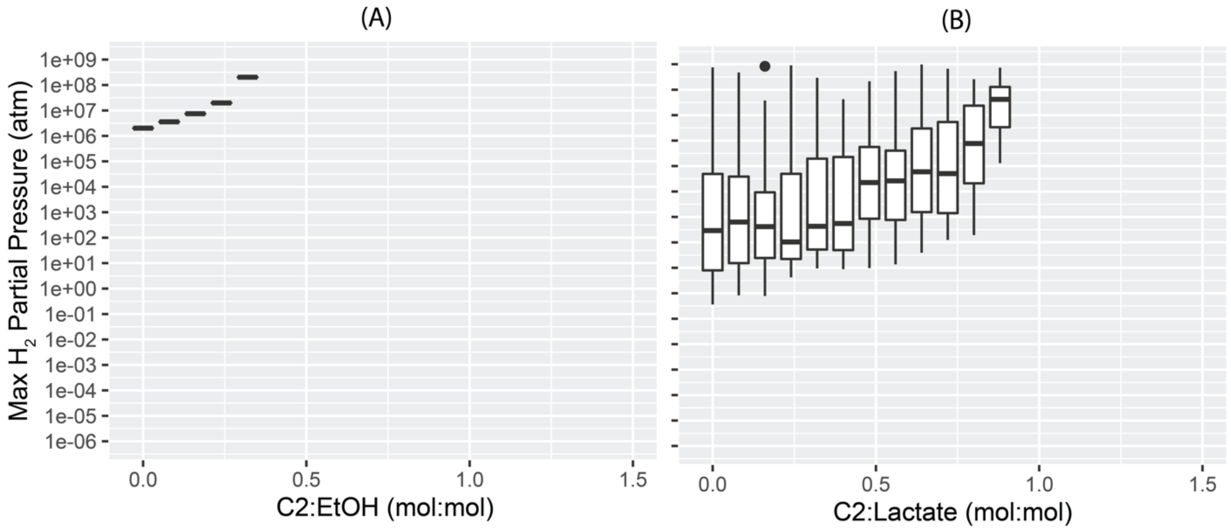


**Figure S7.** Maximum partial pressures for the iFermCell215 results predicted with (A) acetate and ethanol as co-substrates and (B) acetate and lactate as co-substrates. Values greater than $10^9$ atm are not shown. These values correspond to the scenarios plotted in **Fig. 4**.

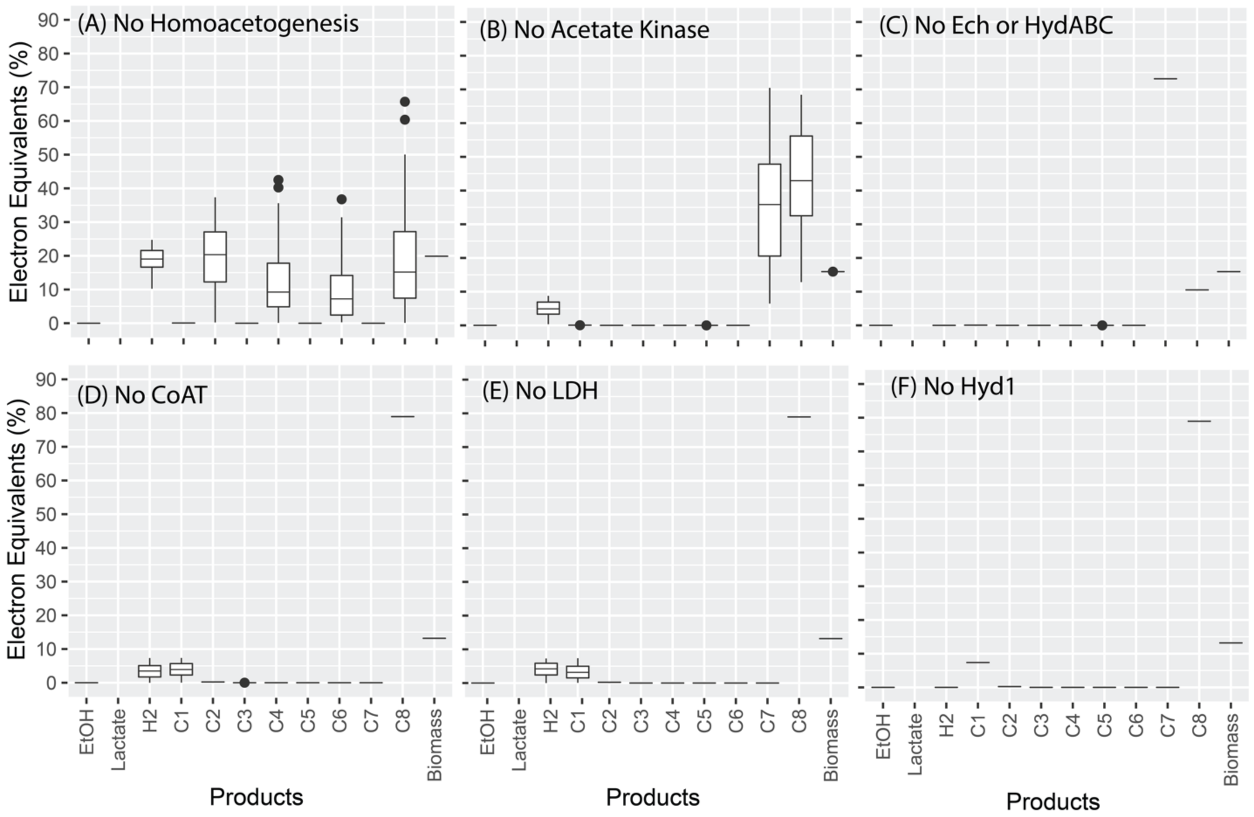


**Figure S8.** Results from additively disabling reactions in iFermCell215 when feeding lactate. (A) disabling homoacetogenesis; (B) disabling homoacetogenesis and acetate kinase; (C) disabling homoacetogenesis, acetate kinase, and energy conserving hydrogenases (Ech and HydABC); (D) disabling homoacetogenesis, acetate kinase, energy conserving hydrogenases, and CoAT; (E) disabling homoacetogenesis, acetate kinase, and energy conserving hydrogenases (Ech and HydABC); (F) disabling homoacetogenesis, acetate kinase, energy conserving hydrogenases, CoAT, and a NADH dependent lactate dehydrogenase (LDH) which requires the model to use an electron-confurcating lactate dehydrogenase reaction; (G) disabling all hydrogen production.

**Table S6.** Calculated eeq contained in substrates and products from a mixed culture fermentation of lignocellulsoic biorefinery residue. Substrates represent the eeq consumed by the microbial community and products represent the millielectron equivalents (meeq) of the measured products. We assumed that the complex carbohydrates remaining in conversion residue were comprised of 50% xylans and 50% glucans, by mass based on published compositions of this material.[38] The bioreactor performance data is from day 96 of reactor operations described in Scarborough, et al 2018.[29]

| **Compound** | **meeq/hr** | **% of eeq** |
|---|---|---|
| **Substrates** | | |
| Glucose | 0.300 | 6.1% |
| Xylose | 2.635 | 53.9% |
| Glucans | 0.780 | 16.0% |
| Xylans | 0.731 | 15.0% |
| Lactate | 0.007 | 0.1% |
| Glycerol | 0.433 | 8.9% |
| **Products** | | |
| Ethanol | 0.112 | 2.3% |
| Formate | 0.000 | 0.0% |
| Acetate | 0.625 | 12.8% |
| Propionate | 0.000 | 0.0% |
| Butyrate | 1.755 | 35.9% |
| Valerate | 0.000 | 0.0% |
| Hexanoate | 1.615 | 33.0% |
| Heptanoate | 0.000 | 0.0% |
| Octanoate | 0.161 | 3.3% |
| $H_2$ | 0.383 | 7.8% |
| Biomass | 0.236 | 4.8% |

**Table S7.** Summary of additional constraints for modeling the bioreactor community as a set of functional guilds. An "X" indicates the model has the indicated capability. Greyed out boxes indicate that the functional guild model does not model this activity.

| | | Carbohydrate Utilization | | | | | | Hydrogenase | | |
|---|---|---|---|---|---|---|---|---|---|---|
| Reactor Population | Functional Guild | Xylans | Glucans | Xylose | Glucose | Odd-chain Production | Acetate Kinase | HYD1 | ECH | HYDABC |
| Cand. *W. Bifida* | SEOs | | | X | X | | X | X | X | |
| Cand. *P. fermentans* | LEOs | | | | | | | X | X | |
| *Lactobacillus* | SFOs | X | X | X | X | | X | | | |
| *Coriobacteriaceae* | HSFs | | X | | X | | X | X | | |

**List of Supplementary Files:**

**Supplementary Data File 1:** Calculations for obtaining the simplified biomass equation

**Supplementary Data File 2:** iFermCell215 Metabolic Model in .py format

**Supplementary Data File 3:** iFermGuilds564 Metabolic Model in .py format

**Supplementary Data File 4:** Lists of reactions and metabolites for both metabolic models

**Supplementary Data File 5:** Flux predictions from metabolic modeling results used in this study
